# The evolution of stomatal traits along the trajectory towards C_4_ photosynthesis

**DOI:** 10.1101/2020.11.10.377317

**Authors:** Yong-Yao Zhao, Mingju Amy Lyu, FenFen Miao, Genyun Chen, Xin-Guang Zhu

## Abstract

C_4_ photosynthesis optimizes plants carbon and water relations, allowing high photosynthetic rate with low stomatal conductance. Stomata have long been believed as a part of C_4_ syndrome. However, it remains unclear how stomata traits evolved along the path from C_3_ to C_4_. Stomatal patterning was examined in *Flaveria* genus, a model for studying C_4_ evolution. Comparative, transgenic and semi-*in-vitro* experiments were used to study molecular basis that underlies stomatal traits along C_4_ evolution. Novel results: the evolution from C_3_ to C_4_ species through intermediate species is accompanied by a stepwise rather than an abrupt change in the stomatal traits. The initial change occurs near Type II and dramatic change occurs at the C_4_-like species. On the road to C_4_, stomata become less in number but bigger in size and changes in stomatal density dominates changes in maximum stomatal conductance (*g_smax_*). The reduction of *FSTOMAGEN* expression underlies altered *g_smax_* between *Flaveria* species with different photosynthetic pathways and likely occurs in other C_4_ lineages. Our study provides insight into the pattern, mechanism and role of stomatal evolution along the road towards C_4_. This work highlights the stomatal traits in the current C_4_ evolutionary model and the co-evolution of photosynthetic pathway and stomata.

## Introduction

Stomata are small pores surrounded by a pair of guard cells on the leaf surface, which control the passage of water and CO_2_ between leaf and atmosphere (*1*). The emergence of C_4_ photosynthesis optimizes plant water and carbon relations, allowing low stomatal conductance (*g_s_*) (i.e reduced water loss) without compromising carbon assimilation (*2–4*), which raises a possibility that C_4_ photosynthesis might have been selected as a water-conserving mechanism (*3*) and that water limitation might be a primary driver of C_4_ evolution (*5–7*). Comparison study on multiple independent lineages of C_4_ grasses showed that the g_s_ is significantly lower in C_4_ species relative to C_3_ relatives (*2, 8*), and the higher intrinsic water use efficiency (iWUE) of C_4_ plants compared to the C_3_ plants can be attributed to both lower *g_s_* and higher photosynthetic CO_2_ uptake rate (*A*) (*2, 9–11*). The high carbon assimilation capacity of C_4_ plants contributes to their high biomass, while low g_s_ is crucial for C_4_ plants to better adapt to saline land, and hot, arid & open habitat (*12, 13*).

In a long-term adaptation, C_3_ plants generally adjust their maximal stomatal conductance (*g_smax_*) (*14–16*). Phylogenetically controlled comparison shows that C_4_ grass have lower *g_smax_* compared with their close related C_3_ grass (*17*). In line with this, C_4_ grass also have lower operating *g_s_* (*2, 17*). Though low *g_smax_* is regarded as an important feature in the C_4_ syndrome (*18*), the changes of *g_smax_* along the road to C_4_ photosynthesis have not been studied (*19*). *G_smax_* is determined by stomatal density (*SD*) and stomatal size (*SS*). Although higher *SD* of C_4_ species compared to C_3_ species has been observed (*20*); in some cases, C_4_ plants can have a lower *SD* compared to C_3_ species (*21*). These differential changes in stomatal traits between photosynthetic types might be related to phylogenetic effects (*2, 22, 23*). As a reflection of this, comparative study using plants from different C_4_ clades shows that lower *g_smax_* in C_4_ plants is due to either a smaller stomatal size (*SS*) for some C_4_ clades, or lower stomatal density (*SD*) for other clades (*17*), but after accounting for the phylogenetic effects, *SD* in C_4_ is lower than C_3_ species and there is no significant difference in *SS* between C_3_ and C_4_ grass (*17*). Earlier study suggests that the influence of photosynthetic pathway on *SD* and *SS* is only statistically marginally significant (*17*), which might because stomatal development is affected by many factors, especially the environmental conditions such as soil moisture and fertility status (*24–27*). Therefore, more evidence is still needed to study the changes in *SS* and *SD* during C_4_ evolution.

Stomatal development is controlled by a well-studied genetic program (*28*), which involves a series of ligands and transmembrane receptors. The ligands and receptors lead to downstream changes of mitogen-activated protein kinase (MAPK) and transcription factors (*28*). TOO MANY MOUTHS (TMM) and ERECTA-family (ERfs: ER, ERL1, ERL2) are the transmembrane receptor-like kinases that negatively regulate stomatal development (*29, 30*), and the TMM interacts with ERfs (*29, 31*). The ligands that bind to ERfs and TMM receptors are mainly EPIDERMAL PATTERNING FACTOR/EPIDERMAL PATTERNING FACTOR-LIKE family (EPF/EPFL-family) peptides (*32*). Overexpression of *EPF1* decreases *SD* (*33*). *EPF2* has a similar genetic effect as *EPF1* (*34*). STOMAGEN prevents the phosphorylation of downstream receptors, which results in the changed expression of basic Helix-loop-Helix (bHLH) paralogs, SPCH, MUTE and FAMA, ultimately positively regulates stomatal development (*35–38*).

C_4_ plants evolved from C_3_ ancestors through a number of intermediate stages, which were shown in a number of genera (*7, 39–41*). Among these genera, we chose *Flaveria* genus as the model for this study. The *Flaveria* genus contains close species with multiple different photosynthetic types which cover the sequential stages of C_4_ evolution, i.e. C_3_, C_3_-C_4_ type I, C_3_-C_4_ type II, C_4_-like and C_4_ (*4, 41–45*). This genus is the youngest genus in which C_4_ photosynthesis evolved (*46*) and have many intermediate species (*47*). A close relationship between *Flaveria* species can minimize the impact of non-photosynthetic effects on stomatal traits (*11, 47*). The species in *Flaveria* genus grows on similar growth habits and also have similar morphologies (*13, 48–51*). These make *Flaveria* genus an ideal system to study the evolutionary process from C_3_ to C_4_ (*39*). The physiology, structure, biochemistry and molecular changes of the *Flaveria* genus along C_4_ evolution have been systematically evaluated (*41–45, 47, 51, 52*). In fact, the current C_4_ evolutionary model illustrating the path from C_3_ to C_4_ photosynthesis is largely based on studies on the *Flaveria* species (*39, 53*). The current model of C_4_ evolution include many steps *(41, 45, 47, 48, 51, 52, 54)*. First, bundle sheath cells and vein density showed C_4_ characteristics (*39, 43, 55*). Then, glycine decarboxylase (GDC) transfers from mesophyll cells to bundle sheath cells (*56*), resulting in a modest CO_2_ concentrating mechanism (*55*), which bridges the formation of C_4_ metabolism (*39, 41*). Although stomata have long been thought as a part of C_4_ syndrome (*17, 18*), the evolutionary changes of stomata traits in current C_4_ evolutionary model have been much less studied.

This study aims to answer the following questions: 1) In the context of the current C_4_ evolution model, will stomatal traits and *g_smax_* change with the evolution of photosynthetic type? How and when do stomatal traits change along the path to C_4_? 2) What’s the molecular mechanism underlying the change in the stomatal traits during C_4_ evolution? 3) What are the roles and drivers of stomatal evolvement in the evolutionary process towards C_4_ photosynthesis?

## Result

### Maximum stomatal conductance and stomatal pattern during the evolution of C_4_ photosynthesis in *Flaveria*

As a model of C_4_ evolution, there are distinct evolutionary phases in *Flaveria* genus (Fig 1a). The maximum value of operational *g_s_* (marked with circles on figure) for C_3_ *Flaveria* species under saturated light intensity was higher than that for C_4_ species, indicating that the C_3_ *Flaveria* species tends to have a higher *g_s_* (Supplementary Fig 1a). The pore length of stomata was approximately equal to 0.41×stomatal length, and the guard cell width was approximately equal to 0.33×stomatal length (Supplementary fig. 19). Through measuring stomatal density and stomatal length, we calculated the *g_smax_* for *Flaveria* species. The results show that, whether growing under outdoor natural light or constant light in the greenhouse, the *g_smax_* decreased gradually from C_3_ through intermediate species to C_4_ species (Fig 1b,c,d,e). A significant correlation was found between the *g_smax_* and *g_s_* under saturated light for the *Flaveria* species with different photosynthetic types (Fig 1f). Under saturated light, the *g_s_* of *son* (Type I) was extremely high (marked with circle on figure), matching with its highest *g_smax_* within the *Flaveria* genus (Fig1d, Supplementary Fig 1a). Similar to that under the saturated light intensity, the WUE (*A*/*g_s_*) of *rob* (C_3_) measured under unsaturated light intensity was higher than that of *bid* (C_4_), and the *g_s_* of *rob* was 71% higher than that of *bid* (Supplementary Fig. 1d).

**Fig 1:**
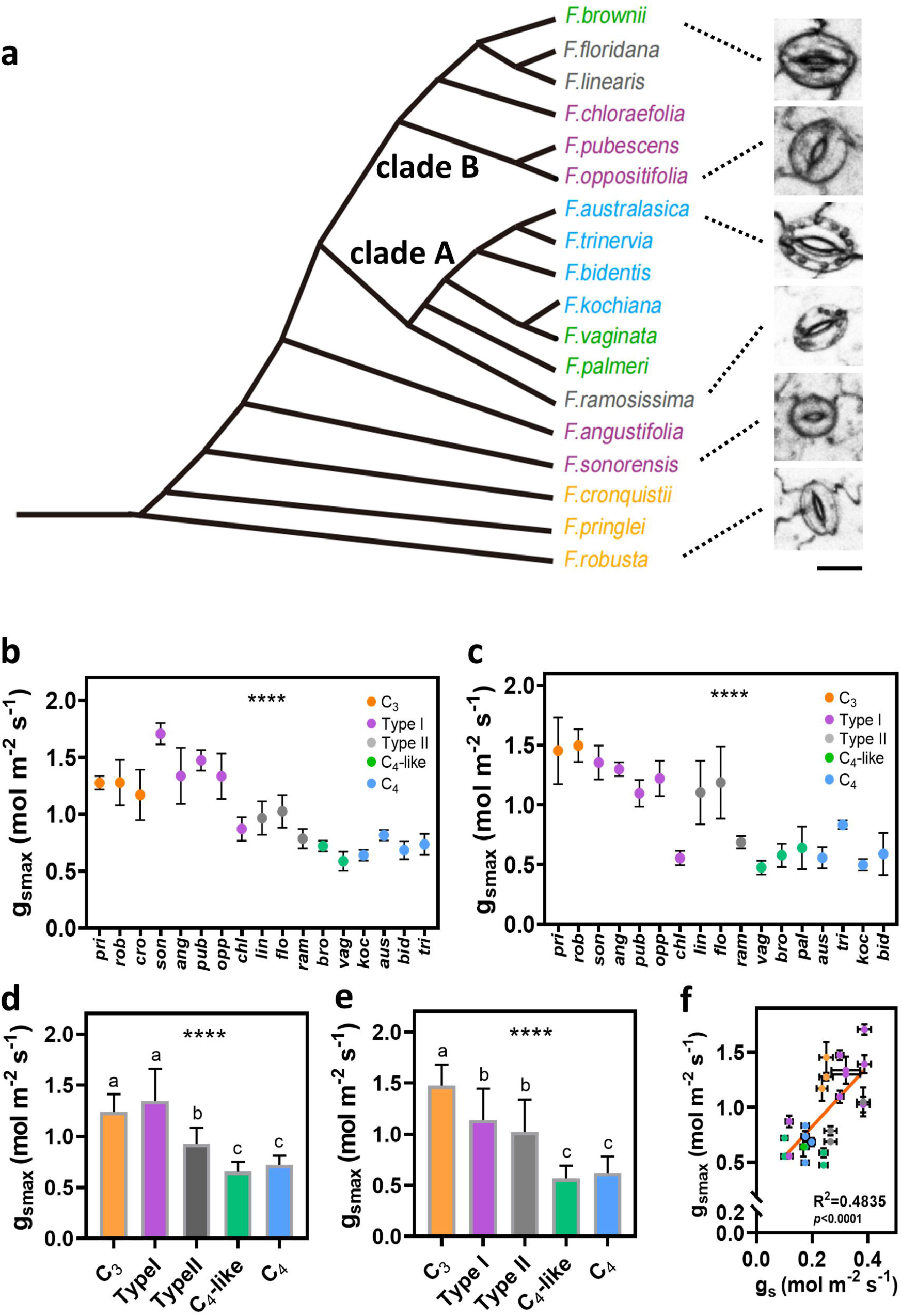
Maximum stomatal conductance (*g_smax_*) of *Flaveria* species **a**, Phylogeny of the Flaveria genus according to (*42, 48*). Different colors represent different photosynthetic types, orange indicates C_3_, purple indicates Type I, gray indicates Type II, green indicates C_4_-like, blue indicates C_4_. **b**,**c**, The trend of *g_smax_* on the abaxial surface in *Flaveria* species with different photosynthetic types from C_3_ to C_4_ grown outdoors (**b**) and in greenhouse (**c**). **d**,**e**, The difference in *g_smax_* of the abaxial surface between species with different photosynthetic types in the *Flaveria* genus grown outdoors (**d**) and in greenhouse (**e**). **f**, Correlation of *g_smax_* and *g_s_* for *Flaveria* species with different photosynthetic types. The data of *gs* was from (*4*). Error bar indicates s.d. The asterisks represent statistically significant differences, One-way ANOVA: *, *P* < 0.05, **, *P* < 0.01, ***, *P* < 0.001, ****, *P* < 0.0001; Different letters represent statistically significant differences, Duncan’s multiple range test (α = 0.05). Pearson correlation coefficient and the significance were noted in each panel.

In comparison to C_4_ and C_4_-like photosynthesis, the stomatal densities (*SD*) of C_3_ and Type I photosynthesis on the abaxial leaf surface of *Flaveria* were nearly 150% higher (Fig. 2a,b,c,d). With the progression from C_3_ to C_4_ species, there was a gradual decrease in *SD* (Fig. 2a, c). The difference in *SD* and *SS* between C_3_ and Type I species wasn’t as large as that between C_3_ and C_4_ species; and the *SD* and *SS* of Type II species fell in between Type I and C_4_-like species (Fig. 2b,d). The *SD* of C_4_ and C_4_-like species were similar, although the *SD* of C_4_-like species was slightly less than that of C_4_ (Fig. 2b,d). For the adaxial leaf surface, the *SD* of C_3_ species was again higher than that of the C_4_ species, although the difference between *SD* in the intermediate species was less significant (Supplementary fig. 3a, b, c, d). There was a strong positive correlation of *g_smax_* and *SD* either on the abaxial leaves (R^2^=0.8955, p<0.0001) (Fig. 2e) or on adaxial leaves (R^2^=0.9165, p<0.0001) (Supplementary fig. 3e) and a significant negative correlation of *g_smax_* and *l* on abaxial leaves (R^2^=0.2596, *p*<0.005) (Fig. 2f), indicating the *SD* determined the change of *g_smax_*, not stomatal size (SS). *SD* and *l* showed a significant negative correlation on abaxial leaves (R^2^=0.5093, *p*<0.0001) (Fig. 2g) or on adaxial leaves (R^2^=0.2055, *p*<0.05) (Supplementary fig. 3g).

**Fig 2:**
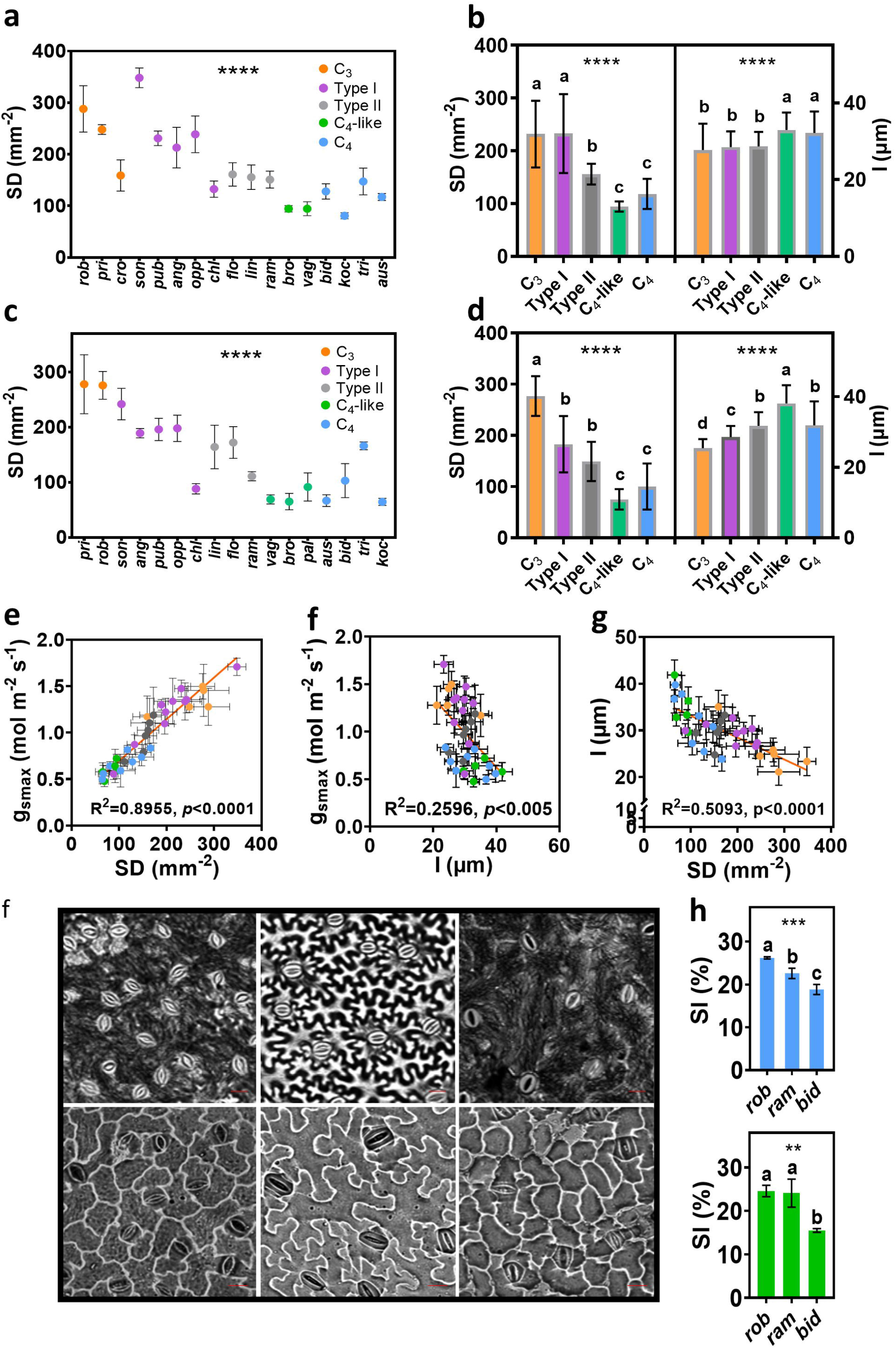
Stomatal density (*SD*) and its patterning. **a**,**c**, The trend of *SD* of abaxial surface from C_3_ to C_4_ in *Flaveria* species grown outdoors (**a**) and in greenhouse (**c**). **b**,**d**, The difference in *SD* and guard cell length (l) of abaxial surface between species with different photosynthetic types in *Flaveria* genus grown outdoors (**b**) and in greenhouse (**d**). **e**, Correlation of *g_smax_* and *SD* for different *Flaveria* species. Different color indicates different photosynthetic type. **f**, Correlation of *g_smax_* and *l* for different *Flaveria* species. Different color indicates different photosynthetic type. **g**, Correlation of *l* and *SD* for different *Flaveria* species. Different color indicates different photosynthetic type. **h**, The photographs of stomatal patterning on abaxial surface in *rob*, *ram* and *bid*. The photographs from the left to right are for *rob*, *ram* and *bid* grown outdoors (top) and in greenhouse (bottom). **i**, Comparison of stomatal index (*SI*) between *rob*, *ram*, and *bid*. Different colours represent plants grown outdoors (blue) or in greenhouse (green). *SD*, (n=4, biologically independent replicates). *l*, (n=20, biologically independent replicates). *SI*, (n=3, biologically independent replicates). Error bar indicates s.d. The asterisks represent statistically significant differences, One-way ANOVA: *, *P* < 0.05, **, *P* < 0.01, ***, *P* < 0.001, ****, *P* < 0.0001; Different letters represent statistically significant differences, Duncan’s multiple range test (α = 0.05). Pearson correlation coefficient and the significance are noted in each panel.

*Flaveria* species, i.e. *rob*, *ram* and *bid*, were chosen to represent C_3_, intermediate and C_4_ species, respectively, to analyse the ratio between stomatal number and the number of epidermal cells, i.e. the stomatal index (*SI*). The *SI* of *rob* was nearly 50% higher than *bid* and *ram* in plants grown in either greenhouse or outdoors, suggesting that the proportion of stomata occupying the epidermis, contributed to the highest *SD* in *rob* (Fig. 2h). In addition to the analysis of *Flaveria* plants grown from cuttings, we also planted *Flaveria* germinated from seeds. Results show that the *SD* of C_3_ species was higher than C_4_ species, consistent with the results from cuttings (Supplementary Fig. 4).

### Comparison of the sequence and transcription abundance of genes related to stomatal pattern between C_3_ and C_4_ *Flaveria* species

The signaling pathway for stomatal development has been studied extensively (*57*). We identified amino acid sequence encoded by homologs of stomatal development related genes in *Flaveria*. Amino acid sequence alignment analysis showed that epidermal patterning factor/epidermal patterning factor-Like gene family (*EPF*/*EPFL*-family) homologs was conserved at the C-terminal part of amino acid sequence (Fig. 4A, Supplementary fig. 5). Members of *EPF*/*EPFL*-family genes encode secretory proteins containing signal peptides at the N-terminus and functional domains at the C-terminus (*33, 34, 58*). Analysis shows that the hydrophobicity of EPFs/EPFL homologs in *Flaveria* was similar to EPF/EPFL in Arabidopsis (Supplementary fig 6), implying the homologs are also the secretory proteins and should have the same physiological function as EPF/EPFL. We hypothesized that either the amino acid sequence or the expression of stomatal development related genes changed during C_4_ evolution. Analysis results reveals no differences within the functional region of amino acid sequence for STOMAGEN, EPF1 and EPF2 homologs between C_3_ and C_4_ species (Fig. 5a, Supplementary fig. 5a, b), indicating the change of amino acid sequence is unlikely to be a cause of the changed *SD* in C_4_ evolution.

As up-regulation of transcription has been a major approach to increase the expression of many genes participating in the building of C_4_ syndrome during C_**3**_ to C_**4**_ evolution in *Flaveria* **(*41, 47*)**, we compared the transcript abundance of stomatal development related genes between C_3_ and C_4_ *Flaveria* species. The expression patterns of known C_4_ related genes in our data analysis were same as the previous works, which indicates that our data collection and analysis procedure were reliable (Supplementary Fig. 11) (*41, 47*). We found that the homologs of most stomatal development related genes showed higher transcript abundance in developing tissues as compared to developed tissues (Fig. 3b), which is consistent with their function of regulating stomatal differentiation in developing leaf (*33, 36, 38, 59*). *STOMAGEN*, *EPF1*, *SDD1*, MPK3 and *TMM* showed obviously lower expression levels in C_4_ species than in C_3_ species (Fig. 3c). Among these genes, *STOMAGEN* was the one that changed the most (Fig. 3c) and showed the most significant difference (Fig. 3d). *EPF1*, *SDD1, MPK3* and *TMM* are negative regulators for stomatal development (Supplementary table 3) (*32–34, 60*). Actually *SD* in C_4_ species was lower than that in C_3_ (Fig 2a,b,c,d), indicating these genes should not be the factors responsible for the decrease in *SD* of C_4_ species. *STOMAGEN* is a positive regulator of *SD* (Supplementary table 3) (*32, 36, 37*), whose lower expression in C_4_ species was in line with the decreased *SD* in C_4_ species (Fig.2a,b,c,d,Supplementary Fig. 3c, d). RT-qPCR quantification of the expression levels of *STOMAGEN*, *EPF1*, *EPF2* homologs between C_3_ and C_4_ species grown under our conditions also confirmed the results from RNA-SEQ (Fig. 3e, Supplementary fig. 14). In this study, we renamed *STOMAGEN* in *Flaveria* as *FSTOMAGEN*.

**Fig 3:**
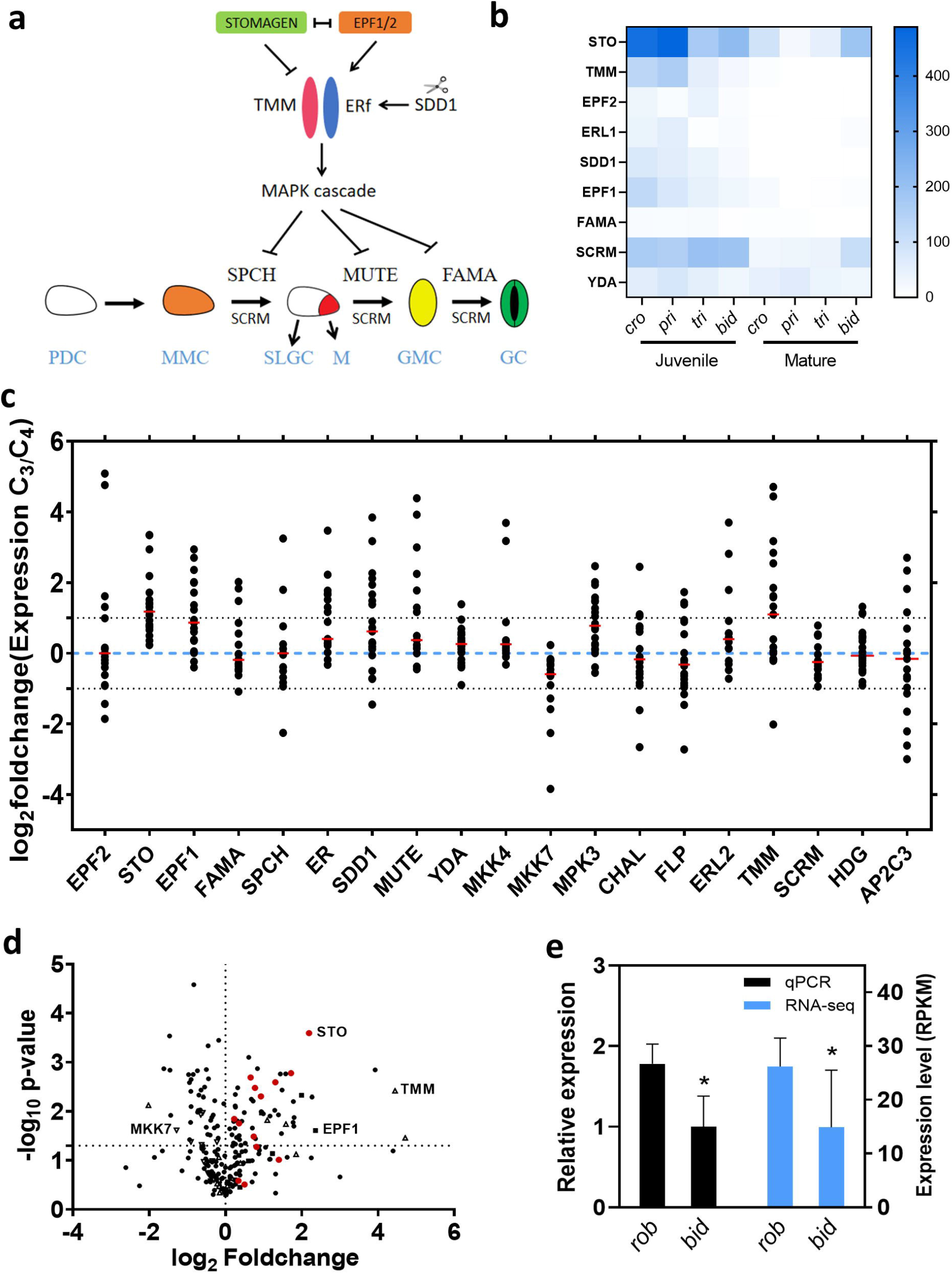
Transcript abundance of stomatal developmental related genes between C_3_ and C_4_ *Flaveria* species. **a**, A model of the signaling pathway for the stomatal development in Arabidopsis (*28*). A protodermal cell (PDC, white) becomes a meristemoid mother cell (MMC, orange), which divides asymmetrically into two daughter cells, with one being a meristemoid (M, red) and the other being a stomatal-lineage ground cell (SLGC). The M cell develops into a guard mother cell (GMC, yellow); the SLGC develops into a pavement cell (PC). The GMC conducts a symmetrical division to produce two equal-sized guard cells (GC, green). EPF1, EPF2 and STOMAGEN competitively bind to ER and TMM, which can deliver a signal to the YDA MAPK cascade. SPCH, MUTE and FAMA ultimately are inactivated through phosphorylation by the YDA MAPK cascade. **b**, Comparison of the expression of stomatal developmental related genes between juvenile and mature leaves. **c**, The foldchanges of stomatal development related genes for the C_3_ and C_4_ *Flaveria* species. Fold change means the ratio between the gene expression in a C_3_ *Flaveria* specie and gene expression in a C_4_ *Flaveria* specie under the same leaf development stage and growth condition as the C_3_ *Flaveria* species. The expression data were from (*66*), (*41*), (*47*) and One Thousand Plants (1KP). **d**, The volcano plot for the expression of the stomatal development related genes between C_3_ and C_4_ *Flaveria* species. The expression data were from (*41*) and (*66*). **e**, Relative expression level determined by RT-qPCR and expression level determined by RNA-seq of *FSTO* (*FSTOMAGEN*) in *rob* and *bid*. (n=4, biologically independent replicates). Error bar indicates s.d. The asterisks represent statistically significant differences (*P*<0.05, t-test, two-tailed distribution, unpaired).

There are considerable differences in amino acid sequence of signal peptide of *STOMAGEN* homologs, especially between species with large evolutionary distance (Fig. 4a Fig. 5a). The functional domain of *STOMAGEN* is however relatively conserved, not only between different *Flaveria* species (Fig. 5a), but also between species that have a large evolutionary distance (Fig. 4a). A phylogenic tree constructed for *STOMAGEN* from different plants suggest *STOMAGEN* homologs of species in the genus *Flaveria* are extremely conserved (Supplementary fig. 18). Experiments with exogenously applied STOMAGEN peptides shows that the STOMAGEN induce increased SD extracellularly (*36, 37, 58, 59*). Previous studies showed that overexpressing rice *STOMAGEN* homologs increases SD in Arabidopsis and RNAi silencing of *STOMAGEN* homologs reduces SD in rice (*61–63*), implying *STOMAGEN* homologs also regulate stomatal development in other species besides Arabidopsis. When plants are grown under conditions that promote or inhibit stomatal development, the expression of *STOMAGEN* is known to increase or decrease, respectively (*24, 59, 64*). A similar modulation of *STOMAGEN* expression levels was observed for *rob* when grown at different light conditions (Supplementary Fig 9).

To confirm that *FSTOMAGEN* can induce stomatal generation and assess the ability of *FSTOMAGEN* to regulate stomatal development, we overexpressed the *FSTOMAGEN* in *Arabidopsis*. Specifically, we recombined the signal peptide of *STOMAGEN* from *Arabidopsis* to the functional region of *FSTOMAGEN* and named the hybrid gene as AFSTO (Fig, 4e). Overexpressing either the AFSTO or the intact *FSTOMAGEN* (FSTO) from *F.rob* into *Arabidopsis* resulted in similar increased *SD* and could exceed 800 stomata per unit leaf area (800 mm^−2^) (Fig. 4f,g,i,j,h, Supplementary fig 15). Identification of transcription showed successful transgenes and the expression level of the AFSTO or FSTO increased with the increase in SD (Fig. 4f, g, i, j). Furthermore, overexpression of *FSTOMAGEN* also induced clustered stomata (Fig. 4k, Supplementary fig 20), as also reported earlier for the over-expression of *STOMAGEN* (*36*). To further confirm that *FSTOMAGEN* can increase *SD* in *Flaveria*, we synthesized a peptide representing the FSTOMAGEN functional region, consisting of 45 amino acids. In order to confirm our synthesized peptides that have been folded were biologically active, the FSTOMAGEN peptides were applied *in vitro* on the leaves of Arabidopsis and resulted in increased *SD* and clustered stomata (Fig 4b, Supplementary fig 17b). Application of the peptides on leaves of *bid* also increased *SD* and *SI* and caused stomatal cluster (Fig 4c, d, Supplementary fig 17a). These results demonstrate that, in addition to its *in vivo* function, *in vitro* application of FSTOMAGEN also directly positively regulate *SD* in *Flaveria*. This is consistent with the notion that *STOMAGEN* is expressed in mesophyll cells and secreted out of cells to influence the stomatal development in the epidermal layer in *Arabidopsis* (*36, 37*).

**Fig 4:**
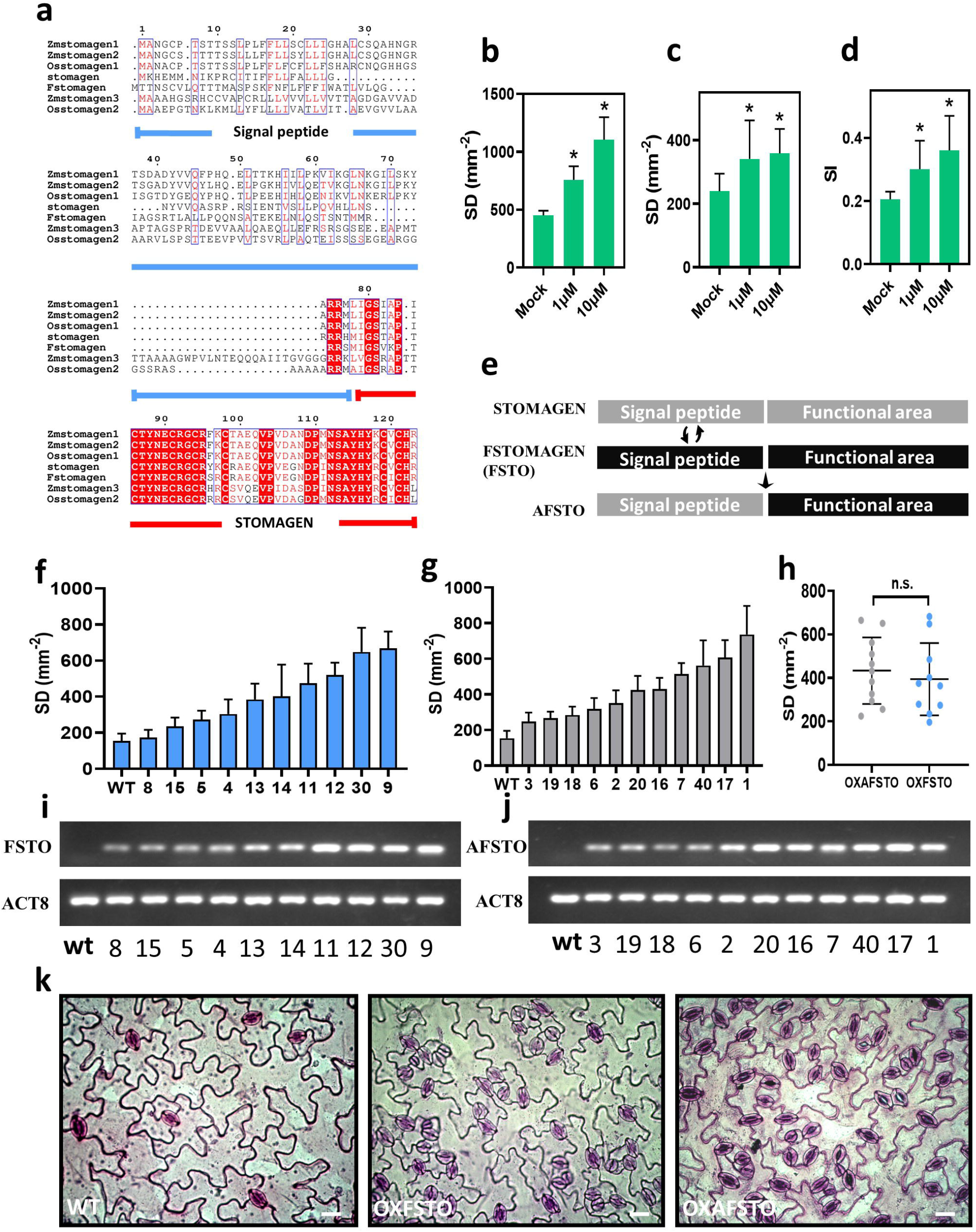
The Function of *FSTOMAGEN* in controlling the *SD* **a**, Protein conservation analysis of the homologs of FSTOMAGEN in different species. The amino acid regions marked by green line and red line represent the signal peptide and the functional domain of FSTOMAGEN, respectively. Red letters represent conserved amino acids in most species, white letters with red backgrounds represent amino acid conserved across all species. **b**, The stomatal density for Arabidopsis leaves with FSTOMAGEN applied at different concentrations (n=8, biologically independent replicates). **c**, The stomatal density for bid leaves with FSTOMAGEN applied at different concentrations (n=8, biologically independent replicates). **d**, The stomatal index (SI) for bid leaves with FSTOMAGEN applied at different concentrations (n=6, biologically independent replicates). Error bar indicates s.d. The asterisks represent statistically significant differences (*P*<0.05, t-test, two-tailed distribution, unpaired). **e**, The structure of recombinant genes used to transform Arabidopsis. AFSTO represents the recombination of the signal peptide of STOMAGEN from Arabidopsis and the functional domain of FSTOMAGEN from *F.robusta*. FSTO represents the intact STOMAGEN from *F.robusta*. **f**, **i**, The *SD* (**e**) and RNA-level (**h**) of the different transgenic Arabidopsis (FSTO) lines. **g**, **j**, The *SD* (**f**) and RNA-level (**i**) of the different transgenic Arabidopsis (AR) lines. **h**, Comparison of the transgenic FSTO lines and AR lines. **k**, Photographs of stomata in a leaf from Arabidopsis with FSTO and AFSTO over-expressed. Scar bar represents 20 um. Error bar indicates s.d. The n.s. represent statistically no significant differences (P>0.05, t-test, two-tailed distribution, unpaired).

**Fig 5:**
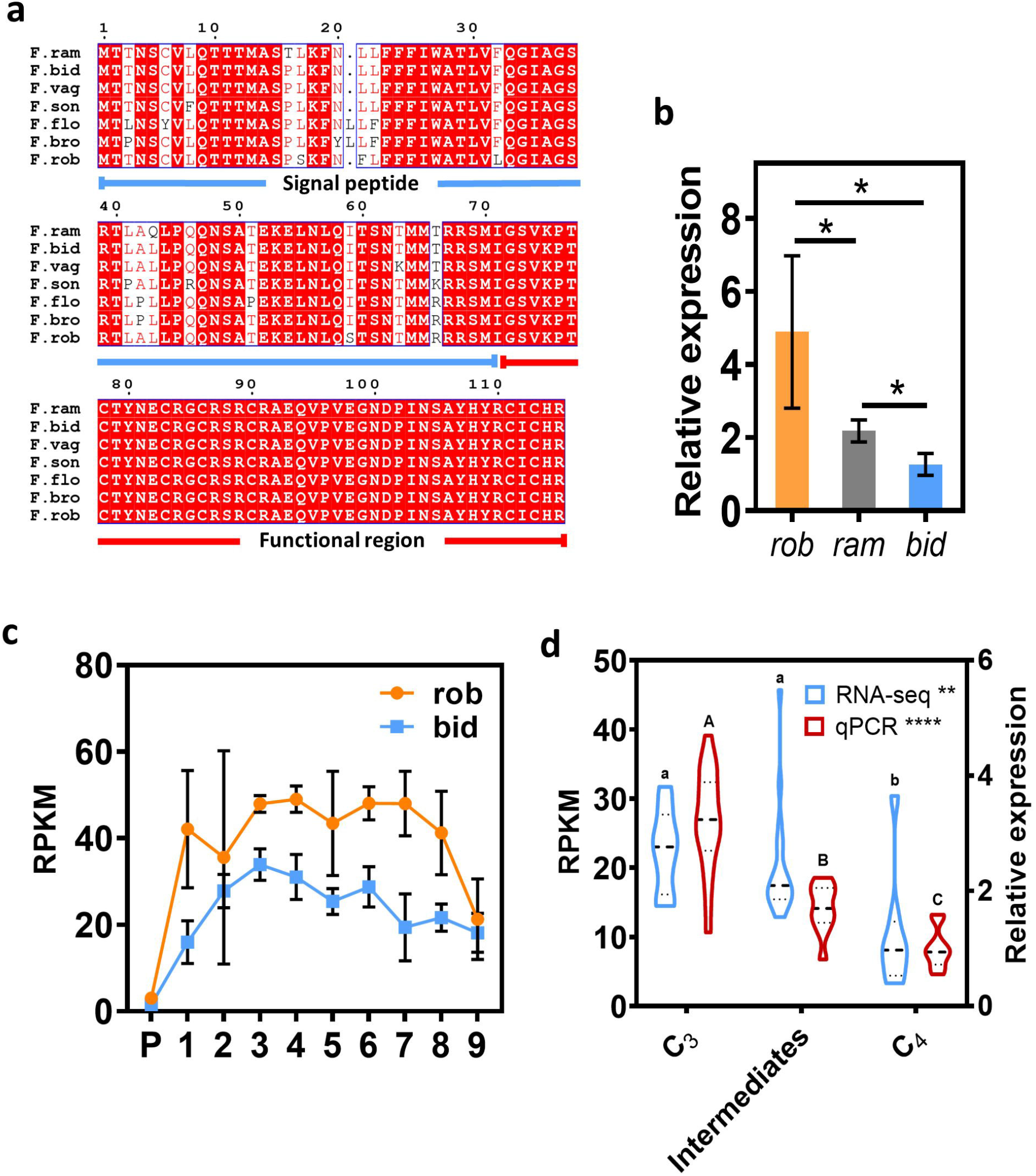
Expression level of *FSTOMAGEN* at different stages of C_4_ evolution. **a**, Protein conservation analysis of the homologs of FSTOMAGEN between different species in the *Flaveria* genus. The amino acid regions marked by green line and red line represent the signal peptide and the functional domain of STOMAGEN, respectively. Red letters represent conserved amino acids in most species; white letters with red background represent amino acids conserved across all species. **b**, Expression levels of *FSTOMAGEN* determined by RNA-seq at the different developmental stages of leaves in *rob* and *bid*. RNA-seq data was from (*66*). P denotes leaf primordia. Approximate leaf length for the different leaf stages was 0.3 cm (stage 1), 0.5 cm (stage 2), 1.5 cm (stage 3), 2 cm (stage 4), 3 cm (stage 5), 4 cm (stage 6), 6 cm (stage7), 8 cm (stage 8) and 10 cm (stage 9). **c**, Comparison of relative expression levels of *FSTOMAGEN* between *rob*, *ram* and *bid*. (n=3, biologically independent replicates). The asterisks represent statistically significant differences (*P*<0.05, t-test, two-tailed distribution, unpaired). **d**, Comparison of the expression levels of *FSTOMAGEN* between different species with different photosynthetic types in *Flaveria* genus. For RT-qPCR, C_3_ species incude *pri* (n=5) and *rob* (n=13), intermediates species includes *ang* (n=3), *ram* (n=5) and *vag* (n=3), C_4_ species include *tri* (n=4), *aus* (n=3), *bid* (n=4) and *koc* (n=3). For RNA-seq, C_3_ species include *pri* (n=4) and *rob* (n=4), intermediates species includes *ano* (n=4), *pub* (n=4), *chl* (n=4), *bro* (n=4), C_4_ speices include *tri* (n=4) and *bid* (n=4). RNA-seq data was from (*41*). The number in parenthesis indicates biologically independent replicates. The relative expression indicates the measurements with RT-qPCR, and the others were measured with RNA-seq. The geometric means of *EF1a* and *ACT7* were used as the reference in the calculation of expression levels for other genes. Error bars indicate s.d. The asterisks represent statistically significant differences, One-way ANOVA: *, *P* < 0.05, **, *P* < 0.01, ***, *P* < 0.001, ****, *P* < 0.0001; Different letters represent statistically significant differences, Duncan’ s multiple range test (α = 0.05)).

### The transcript abundance of *FSTOMAGEN* during C_4_ evolution in *Flaveria*

We examined the copy number of *FSTOMAGEN* in *Flaveria* using blastn with the coding sequence of *FSTOMAGEN* as query sequence against the assembled contigs of each *Flaveria* species from 1KP. Results show that there was only one hit sequence with e-value less than 10^−5^ in each species, indicating that there is a single copy of *FSTOMAGEN* in *Flaveria* (Supplementary fig. 12). Therefore we can ignore the impact of other *FSTOMAGEN* paralogs on stomatal development, and there will be no quantitative errors in the gene expression levels caused by other paralogs that were not considered. Examination of the RNA-seq data of *Flaveria* shows that the expression levels of *FSTOMAGEN* in C_3_ species were markedly higher than those in C_4_ species along different developmental stages of leaves (Fig. 5c). During leaf growth and development, the expression of *FSTOMAGEN* in both *bid* and *rob* increased first, reaching the highest level at leaf development stage 3 (leaf length 1.5 cm) or 4 (leaf length 2.0 cm), and then declined (Fig. 5c). The similar patterns of *FSTOMAGEN* expression in these two *Flaveria* species indicates that it would be valid to compare the *FSTOMAGEN* expression levels in leaves at the same development stage between *Flaveria* species. The expression level of *FSTOMAGEN* in *bid* (C_4_) was lower than that in *rob* (C_3_), and that of *ram* (intermediate species) lied between them (Fig. 5b). This encouraged us to test more *Flaveria* species with different photosynthetic types. Similar results were obtained with either RNA-seq or RT-qPCR (Fig. 5d, Supplementary Fig. 13), where we found the gradual decrease of *FSTOMAGEN* expression from C_3_ to intermediate species to C_4_ species.

### The expression of *FSTOMAGEN* homologs for species with different photosynthetic type in multiple C_4_ lineages

To gain insight into whether *STOMAGEN* homolog might be a generic factor leading to the difference in *SD* between C_3_ and C_4_ species in higher plants, we tested its expression level in different C_4_ lineages other than the *Flaveria genus*. The *SD* of abaxial leaf in C_3_ rice was 240% higher than that in C_4_ maize, consistent with a 570% higher *g_s_* in rice compared to maize (Fig. 6a, c). The stomatal length (*l*) in maize was 133% longer (Fig. 6b). We analysed the transcript abundance of *STOMAGEN* homologs, since there are two and three paralogs of *STOMAGEN* in rice and maize respectively (*61*), only one in rice and two in maize are expressed. The *STOMAGEN* homologs in rice had much higher transcript abundance than those in maize (Fig. 6f), while the transcript abundance of other stomatal development related genes did not differ to the same extent between rice and maize (Fig. 6e). This result encouraged us to test more C_4_ lineages (Fig. 6g). C_4_ photosynthesis related genes were indeed higher in C_4_ species, suggesting the data and analysis pipeline can be used to compare the differences in expression of genes between C_3_ and C_4_ photosynthetic types (Fig 6h). It was showed that the expression levels of *STOMAGEN* homologs for C_4_ were lower than those for C_3_, while intermediate had expression between them (Fig 6i).

**Fig 6.**
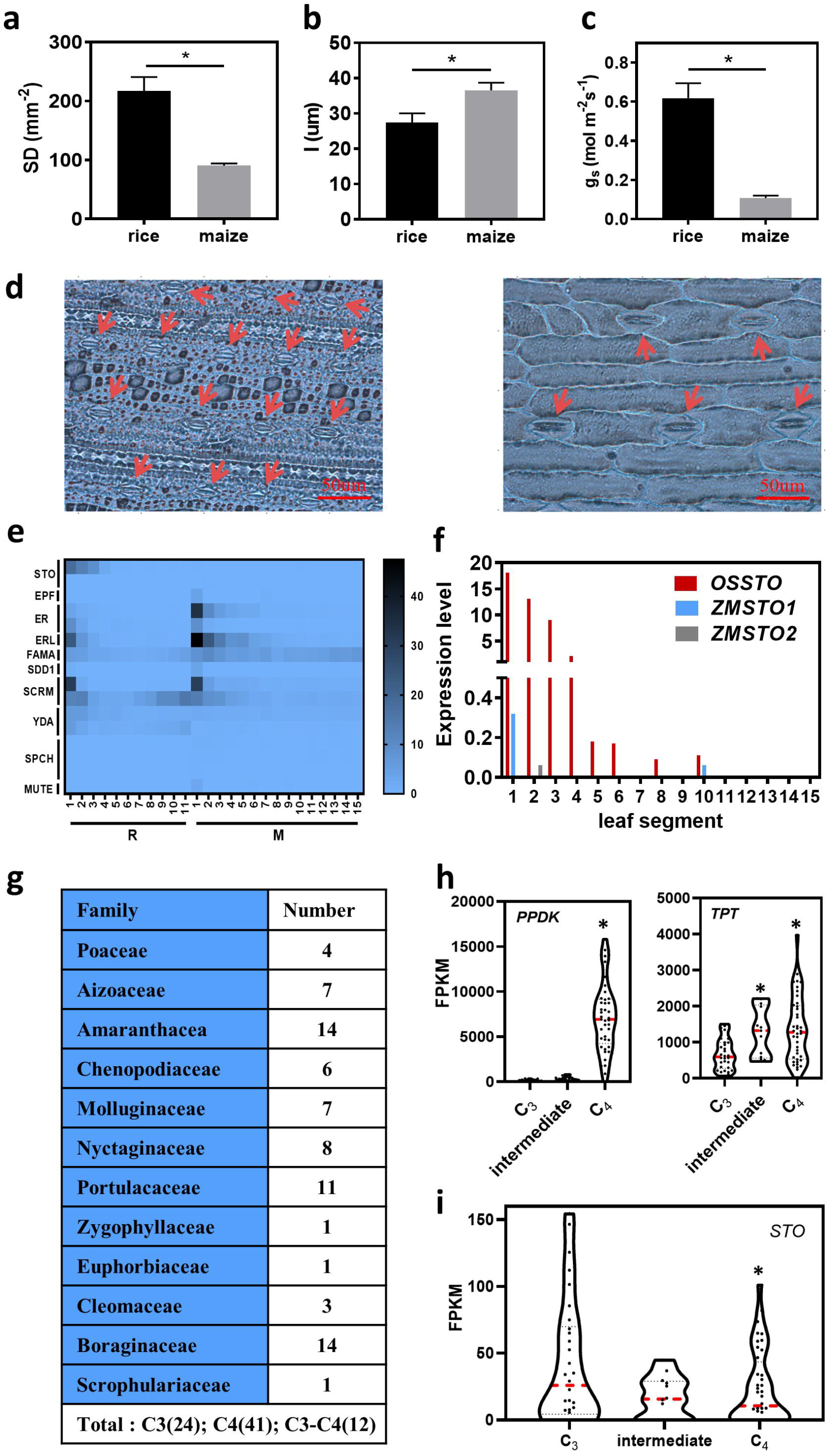
Comparison of stomatal pattern and expression levels of stomatal development related genes in the C_4_ lineages other than *Flaveria* **a**, Stomatal characteristics for rice and maize. *SD*, (n=8, biologically independent replicates); *l*, (n=25, biologically independent replicates); *SI*, (n=3, biologically independent replicates); *g_s_*, (n=8, biologically independent replicates). Error bar indicates s.d. The asterisks represent statistically significant differences (*P*<0.05, t-test, two-tailed distribution, unpaired). **b**, Photographs of stomatal pattern in rice and maize. Scale bars represent 50 μm. **c**, Comparison of transcript abundance of stomatal development related genes between rice and maize. The developmental stages of leaves were divided into 11 and 15 parts in rice and maize, respectively. R represents the leaves of rice, M represents the leaves of maize. **d**, Comparison of the expression levels of *STOMAGEN* homologs between rice and maize. There are one and two paralogs of *STOMAGEN* that is expressing in rice and maize, respectively. *OsSTO* represents the homologs of *STOMAGEN* in rice. *ZmSTO* represents the homologs of *STOMAGEN* in maize. **e**, The families and numbers of species with different photosynthetic type that we used to analyse. **f**, Comparison of the expression of C_4_ related genes between species with different photosynthetic types. **g**, Comparison of the expression of *STOMAGEN* homologs between species with different photosynthetic types. The expression was determined by RNA-seq in rice and maize. The RNA-seq data was from (*69*). The asterisks represent statistically significant differences (P<0.05, t-test, two-tailed distribution, unpaired).

**Fig 7:**
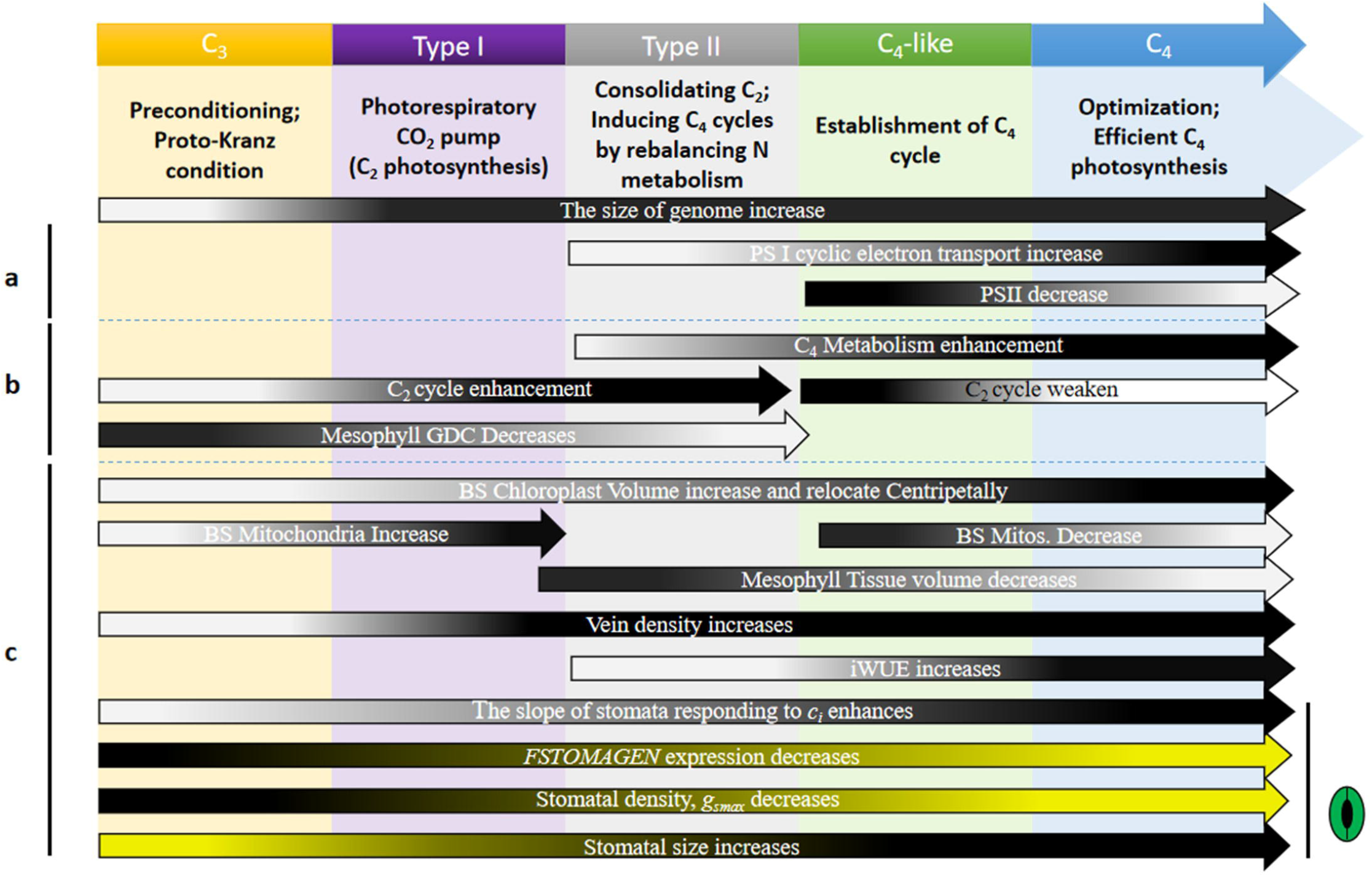
Schematic of the tactic change in the C_4_ syndrome in the current C_4_ model. The position of the C_4_ syndromes in the current C_4_ model was shown. The arrows represent the gradual enhancement or weaken of each feature over the entire evolutionary gradient. a represents photosystem. b represents biochemistry and metabolism. c represents structure and hydraulic feature. Darker color means enhancement, lighter color means weaken. The yellow parts are related to the stomatal traits shown by our data. The rest is the previous research on the C_4_ related features in *Flaveria genus (4, 41, 43–45, 51, 54, 55, 102)*.

## Method

### Plant materials, sample condition and data sources

All *Flaveria* species used in this study were kindly provided by Prof. Rowan F. Sage (University of Toronto). Plants used for stomatal phenotyping were re-grown from cuttings and placed in an environment humidified with humidifiers to generate roots. After plants generated roots, they were transferred outdoors (Institute of Plant Physiology and Ecology, Chinese Academy of Sciences, Shanghai, China), and transplanted into 4-L pots filled with topsoil in April. In July, the number of stomata on abaxial and adaxial surfaces of leaves were counted. Another batch of *Flaveria*, which were also grown from cuttings, were transferred to the phytotron (Center of Excellence for Molecular Plant Sciences, Chinese Academy of Sciences, Shanghai, China). Growth temperature of the phytotron was maintained at 25-28◻, photoperiod was maintained at 16h light / 8h dark, relative humidity (RH) was at 60-65% and photosynthetic photon flux density (PPFD) was maintained at about 180 μmol m^−2^ s^−1^. Two months later, the numbers of stomata on the abaxial and adaxial surfaces of leaves were counted. RNA was extracted from 1.5~3.0 cm long young leaves on the axial shoot which were still expanding. At three hours before sampling, we provided sufficient water to ensure that plants were not water limited, since drought and water status can affect expression levels of *STOMAGEN* (*24, 26*). Gene expression was also examined under normal growth conditions by RT-qPCR for the *rob*, *ram* and *bid*, which are representative of C_3_, intermediate and C_4_ species in the *Flaveria* genus. These three species were grown in the same phytotron with the same environmental conditions, except that we did not water plants before sampling. To test whether plants grown from cuttings or from seeds show difference in the *SD*, we further grew *Flaveria* species from seeds in phytotron and examined stomatal properties. Same environmental conditions were maintained for phytotron as above. Two months after the germination of seeds, we counted stomatal number.

All plants were well watered to avoid drought stress during their growth. We supplied the same amount of commercial nutrient solution (Peters Professional, SCOTTS, N:P:K = 20:20:20+TE, at recommended dose for application) to the different species to avoid nutrient deficiency.

Rice (*Oryza Sativa Nipponbare*) and maize (*Zea maize* B73) were grown in the greenhouse (Center of Excellence for Molecular Plant Sciences, Chinese Academy of Sciences, Shanghai, China) under a PPFD of 550 umol/m^2^/sec, a photoperiod of 12h light / 12h dark, and a daytime temperature of around 31 °C and a night-time temperature of around 22°C. The relative humidity was maintained at around 50-55%. The plants were regularly watered to avoid drought stress.

In addition to the expression data collected by in this study, we also used public data for comparison of transcript abundance. Considering that stomatal development related genes are mainly expressed in young tissues, in each of these used public data source, samples for young leaves were included. For the *Flaveria*, data from four sources for leaves of plants grown under normal condition were analysed. The first data source is the One Thousand Plants (1KP) Consortium, and we subsequently analyzed the data according to the methods described in previous work (*42, 65*). The second and third data sources were Mallman et al (2014) (*41*) and Udo Gowik et al (*47*). The fourth data source was Kumari Billakurthi et al (2018) (*66*). The last three data sources are all from Heinrich Heine University. For the second and third data source, the second and fourth visible leaves were sampled; the second leaf was not completely mature, and stomatal developmental related genes were still actively expressed. In addition to *Flaveria*, transcripts of C_3_ and C_4_ species from other C_4_ lineage sequenced from 1KP were assembled with Trinity (*67*) with default parameters. Transcript abundances were analyzed by mapping the reads to assembled contigs for corresponding samples using the RSEM package with mapping option of bowtie 1 (*68*), and outliers that less than Q1 (lower quartile) − 1.5IQR (interquartile range) and greater than Q3 (upper quartile) + 1.5IQR (interquartile range) were removed. Expression levels for stomatal development related genes in developing leaves for rice and maize were obtained from Wang et al. (*69*).

### Conservation and hydrophobicity analysis of proteins

Amino acid sequence of the homologs of *STOMAGEN*, *EPF1* and *EPF2* were aligned by clustalx, and graphs for the multiple alignments were drawn with ESPrit 3.0 (*70*) (http://espript.ibcp.fr/ESPript/cgi-bin/ESPript.cgi). Hydrophobicity of proteins was determined by ProtScale (*71*) (https://web.expasy.org/protscale/).

### Construction of the phylogenetic tree of *STOMAGEN*

In order to construct the gene evolutionary tree of *STOMAGEN*, we searched the orthologs of *STOMAGEN* in 28 representative species among Viridisplantae as demonstrated in (*72*). The protein sequences of representative species were downloaded from Phytozome (https://phytozome.jgi.doe.gov/). Orthologs of STOMAGEN were predicted applying OrthoFinder (V 0.7.1) (*73*) with default parameters. Proteins sequences of STOMAGEN orthologs were then aligned with MUSCLE (*74*). We then inferred the gene phylogeny using both maximum likelihood (ML) and Bayesian-inference (BI) methods. The ML tree was constructed with RAxML (V2.2.3) (*75*), where bootstrap scores were obtained by running 100,000 independent trees. BI tree was built with MrBayes package (v 3.2.1) (*76*), where posterior probabilities were calculated by running 2000,000 generations with the first 2000 generations being used as burn-in. In both cases, PROTGAMMAILG model (General Time Reversible amino acid substitution model with assumption that variations in sites follow gamma distribution) was used to infer the gene phylogeny. The protein sequence of AT4G14723 (*CLL2*) from *A. thaliana* was used as outgroup as it showed the highest identity with STOMAGEN in *A. thaliana*, with a protein identity of 26.97% and E-value of 0.037 based on Blastp (*77*). Protein sequences for the *Flaveria* species were predicted using OrfPredictor (*78*) based on de novo assembled transcripts as described in (*79*).

### Measurement of stomatal conductance (*g_s_*)

The LI-6400 Portable Photosynthesis System (Li-Cor, Inc., Lincoln, NE, USA) was used to measure *g_s_* on the youngest fully expanded leaves from approximately three-months-old *Flaveria* plants. The cuvette temperature was maintained at 25◻, the PPFD was at ~500 μmol/m^2^s and the CO_2_ concentration was at around 400 ppm. Chamber humidity was matched to ambient air humidity. Leaf-to-air vapor pressure difference (VPD) was maintained around 1.1 kPa. Before the *g_s_* measurements, *Flaveria* plants were maintained under constant light with a PPFD of ~500 μmol/m^2^s until *g_s_* reached a steady state. External environments during the *g_s_* measurements for rice and maize were controlled to be their growth conditions. In maize and rice, the16-day-old third leaves which were the youngest mature leaves, were used for gas exchange measurements (*69*).

### Phenotyping stomatal traits and the calculation of g_smax_

The youngest fully expanded leaves (usually the third or fourth leaves from the apex in *Flaveria*) at the main stem were used to phenotype stomata related traits. In *Flaveria* and maize, stomata related traits were observed using both nail polish method and epidermis directly peeled from leaves (*6, 80*). For the first method, nail polish was painted on both leaf surfaces to produce the impressions of epidermis. After the nail polish was dried, the nail polish was peeled off the leaf and placed on a microscope slide for examination. For the second method, leaf epidermis was peeled off manually and transferred onto microscope slide. For the count of SD in Arabidopsis, the rosette leaves was soaked in 9:1 (V/V) ethanol:acetic acid overnight, washed with water and cleared in chloral hydrate solution (glycerol:chloral hydrate:water, 1:8:1) overnight, and then stomata of Arabidopsis were stained with Safranin-O, finally leaves were placed on a microscope slide for imaging. Light microscope (smarte; CNOPTEC), camera and image processed software (OPTPRO; CNOPTEC) were used to observe and image epidermis at 100x magnification in *Flaveria*. Center of a leaf was chosen to count stomatal density and measure stomatal length. Five stomata randomly chosen along a diagonal of the varnish peels or epidermis were measured for stomatal length.

G_smax_ was calculated with the follow equation (*16*):

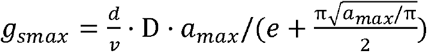

The constants *d* is the diffusivity of H_2_O in air; the constant *v* is the molar volume of air; D is the stomatal density; *a_max_* is the maximum stomatal pore area which is calculated as *a_max_* = *πp*^2^/4 in *Flaveria* species, and the *p* is the stomatal pore length; *e* is the stomatal pore depth, which is known as equal to guard cell width.

### RNA isolation, semiquantitative RT-PCR and quantitative real-time PCR

The young leaf samples taken from the main stem in *Flaveria* or the developing rosette leaves in Arabidopsis were immediately frozen in liquid nitrogen and used directly for RNA extraction. Total RNA was extract from the young leaves using the PureLink RNA Mini kit (Thermo Fisher Scientifc). The first-strand cDNA was synthesized by the TransScript One-step gDNA Removal and cDNA Synthesis SuperMix (TRANSGEN). Semiquantitative RT-PCR was performed with Phanta® Max Super-Fidelity DNA Polymerase (vazyme). The actin8 was chosen as the loading control. Real-time PCR was performed on the CFX connect™ (Bio-Rad) with the UNICON™ qPCR SYBR ®Green Master Mix (YEASEN) according to the manufacturer’s instructions. The calculation of expression according to the 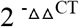.

The expression stability of internal reference genes for the quantitative real-time PCR was assessed by the software: geNORM and NormFinder (*81, 82*). Specifically, we first calculated the coefficient of variations (CV) of all genes in different *Flaveria* species. After that, all genes were sorted according to the calculated CV values. 150 genes with the lowest CV were selected as potential candidate reference genes. Among these, those with mean expression level >300 were chosen as candidate reference genes. We also included those commonly used house-keeping genes, which include *UBC9* (AT4G27960), *UBQ11* (AT4G05050), *ACT7* (AT5G09810), *GAPDH* (AT1G13440), *EF1a* (AT5G60390), *CACMS* (AT5G46630) (Supplementary fig. 6a, b, d), as the internal reference genes in this study. The specificity of the primers (Supplementary table 2) for all these candidate reference genes were identified by agarose electrophoresis (Supplementary fig. 6c). With Genorm (Supplementary fig. 6e) and Normfinder (Supplementary fig. 6f), we concluded that *ACT7*, *EF1a* and *HEME2* showed stable expression levels across *rob*, *ram* and *bid*. *EF1a* and *ACT7* are classic reference genes, and they showed minor variations under different conditions. The geometrical means of the expression level of *ACT7* and *EF1a* were finally used as normalization factors. All primers used for RT-qPCR are listed in Supplementary table 2. For each gene, three technical replicates and at least three biological replicates were performed.

### Vector construction and transformation

A list of primer sequences were used for plasmid construction (Supplementary table 2). The *STOMAGEN* and *FSTOMAGEN* DNA sequences, named ATSTO-cDNA and FloSTO-cDNA respectively, were PCR-amplified from cDNA in Arabidopsis and *rob*. To obtain the AFSTO fragment, the signal peptide region of *STOMAGEN* was cloned from ATSTO-cDNA by PCR-amplification, and the functional region of *FSTOMAGEN* was PCR-amplified from FloSTO-cDNA. The signal peptide and functional region were combined by fusion PCR. To obtain an intact *FSTOMAGEN* (FSTO) gene, the whole gene in *rob*, including the signal peptide, propeptide and the *STOMAGEN* functional region, was amplified from the cDNA of *rob*. To over-express AFSTO and FSTO, the AFSTO and FSTO were cloned separately to a pcambia-3300 binary vector which has a CaMV 35S promoter. This construct was introduced into *A. tumefaciens* strain GV3101, transformed into Arabidopsis using a floral dipping procedure. Transgenic lines were selected on soil that had been treated with a diluted solution of phosphinothricin (diluted 3000 times) over night. About a month after the seeds were sowed on the soil, the stomata were observed on the biggest mature rosette leaf. The phenotype of Arabidopsis T1 and T2 generations was observed at the 400x magnification using the light microscope. For each epidermal peel, five fields were counted for density.

### Refolding of synthetic *FSTOMAGEN*

Chemically synthesized peptide (Synthesized peptide sequence: IGSVKPTCTY NECRGCRSRCRAEQVPVEGNDPINSAYHYRCICHR, Sangon Biotech, Shanghai, China) was dissolved in 20 mM Tris-HCl, pH 8.8, and 50 mM NaCl and then dialysed (Sangon Biotech,0.5 KD-1 KD) for 1 day at 4◻ against 0.5 mM glutathione (reduced and oxidized forms, Macklin) and 200 mML-arginine (meilunbio) at pH 8.0, which were further dialysed three times against 20 mM Tris-HCl, pH 8.8, and 50 mM NaCl for 1.5 days to remove glutathione.

### Stomata induction assay

When the first true leaf or cotyledons appeared in *bid* and Arabidopsis that had germinated in 1/2 MS sterilized solid medium, F*STOMAGEN* peptide was applied on the plants. After the *bid* and Arabidopsis were further incubated in 1/2 MS liquid medium at 22°C for 5 days, stomatal numbers on the abaxial surfaces for the first true leaf and cotyledons were counted under a differential interference contrast microscope (DIC).

### Plotting and statistical analysis

The correlation, linear regressions and t-test (a=0.05) were performed by GraphPad Prism 8. Differences between means of multi-groups were assessed with one-way analysis of variance (ANOVA). Duncan’s multiple range test was performed using the agricolae package of the R (https://www.r-project.org/) to detect differences between two groups. The *A* and *g_s_* at the saturated light intensity were digitized from figure 1 and figure 3 in (*4, 83*) using the OriginPro 2018.

### Accession numbers

The *Flaveria* read data have been submitted to the National Center for Biotechnology Information Short Read Archive under accession numbers: SRP036880, SRP036881, SRP036883, SRP036884, SRP036885, SRP037526, SRP037527, SRP037528, SRP037529, SRP006166, SRP169568. The *Flaveria* RNA-Seq datasets from 1KP Consortium are available at National Center for Biotechnology Information (NCBI) Gene Expression Omnibus (GEO) under accession number: GSE54339. The rice and maize RNA-seq datasets under accession number: GSE54274. Accession numbers of genes in this study was shown in Supplementary table 3.

## Discussion

### Detailed characterization of stomatal traits in the *Flaveria* genus along C_4_ evolution

Many recent studies have shown that stomatal change may play an important role in the evolution of C_4_ photosynthesis (*2, 3, 17*). The short-term regulation of stomata function mainly involves the dynamic opening and closing (*84*). But on an evolutionary scale, the adjustment of stomatal function mainly involves changes in *g_smax_* i.e. changes in stomatal density and stomatal size (*14–17*). The g_*smax*_ in *Flaveria* is significantly correlated with the operating *g_s_* (Supplementary fig 1b). In order to study the stomatal evolution along C_4_ evolution, we used the *Flaveria* genus, which includes species with different photosynthetic pathways. Species in *Flaveria* also has a close evolutionary distance, which can reduce the effects of factors other than photosynthetic types on stomatal traits. In addition, *Flaveria* represents a C_4_ evolutionary model that minimizes differences in habitat (6, 52, 53). The short phylogenetic distance and similar growth habitats enabled us to better detect the difference in stomatal traits between species with different photosynthetic pathways (Fig 1, Fig 2). With data from species in the *Flaveria* genes, we consolidated the earlier notion the *g_smax_* is associated with photosynthetic pathway and there is a decrease in *g_smax_* and *SD* in C_4_ plants compared to C_3_ plants (*17*). Our data further showed that the decrease in *g_smax_* and *SD* also occurred in dicotyledon and this decrease in *SD* is due to less stomata generation (Fig 2h). Photosynthesis and stomata are closely related (*85, 86*), our work provide an excellent example for the co-evolution between photosynthetic pathway and stomata.

We showed that the changes of stomata traits along the path to C_4_ is gradual, not abrupt (Fig 1, Fig 2). The stomata traits initially changed near the Type II stage, and dramatically changed at the C_4_-like stage (Fig 1, Fig 2). Previous studies have shown that in C_4_ grasses the decrease in *SD* might be due to the restriction of stomatal development caused by the increased vein density (*17, 19*). In this study, we found no significant change in stomatal traits in early stage in *Flaveria* (Fig 1, Fig2), while the vein density at this stage, *e.g*. in *F. sonorensis*, was already increased (*43*). Therefore, the decreased *SD* in C_4_ evolution was not necessarily due to the constrained vein development.

Along the road to C_4_, stomata become less in number (lower SD, SI) and larger in size (longer L) (Fig 2g). The opposite trends in *SD* and *SS* changes have been repeatedly shown in different conditions (*14, 16*). The opposite trend may help maintain relative constant allocation of epidermal space to stomata (*14, 16*). Transgenic experiments of genes related to stomatal development have shown that *SS* and *SD* are related, showing that changes in *SD* can directly cause opposite changes in *SS* (*80, 87*). Therefore, the opposite trends of changes in *SD* and *SS* could be caused by the stomatal development related genes, which is consistent with the changes in the expression of *FSTOMAGEN* (Fig 5 b,c,d). Previous studies have shown that the *SS* is related to the size of genome (*88*). Since the genome sizes increased gradually along C_4_ evolution in *Flaveria* (54), the increased genome size might be one reason for the increased *SS* during C_4_ evolution in this genus.

### The role of decreased *g_smax_* during C_4_ photosynthesis evolution in *Flaveria*

Though water limitation might have been regarded as a primary driver for the C_4_ evolution (*5–7*), data from this study suggests that CO_2_ might have been the initial driver for C_4_ evolution. In the *F. sonorensis* and *F. angustifolia*, two Type I intermediate *Flaveria* species, *g_smax_* were still similar to those in C_3_ *Flaveria* species, i.e. *F. cronquistii*, *F. pringlei* and *F.robusta* (*4*) (Fig. 1b,c,d,e) (*89*); while in contrast, the CO_2_ concentration points of these Type I species were already dramatically decreased (*90*) compared to their C_3_ ancestors, as a result of an operating CO_2_ concentrating mechanism supported by the photorespiration CO_2_ pump (C_2_ photosynthesis) (*91*). This is consistent with the notion that at the early stage of C_4_ evolution, where the movement of glycine decarboxylase (GDC) from mesophyll cell to bundle sheath cell occurred, the primary evolutionary driver should be to elevate CO_2_ levels around Rubisco, rather than saving H_2_O (*92*), i.e. the carbon uptake, not water loss, was the major selection pressure at the early stage of C_4_ evolution in this genus (*4, 83*). Only after this initial step, reducing water loss might become crucial. This is because the enhancement of photosynthetic capacity by C_2_ or C_4_ photosynthesis can increase CO_2_ uptake and evapotranspiration of canopies(*4*), hence creating a higher water demand and raising the danger of embolism, especially in the arid, semi-arid, hot and saline areas (*3*). Indeed, we found initial decline in *g_smax_* occurred near Type II (Fig 1b, c, d, e).

The gradually decline of *g_smax_* is consistent with the gradual establishment of photorespiratory C_2_ pump and the shifting of photosynthetic facilities from mesophyll tissue to bundle sheath tissue. The decrease of *g_smax_* represents the negative regulation on intercellular CO_2_ concentration. The higher intercellular CO_2_ concentration could diminish the role of photorespiration CO_2_ pump and photosynthetic capacity in bundle sheath. Indeed, under higher CO_2_, intermediate species will have no photosynthetic advantage over C_3_ plants (*89, 93*). Therefore, it is possible that the gradual decrease of *g_smax_* can help consolidate and strengthen C_2_ photosynthesis during C_4_ evolution. The photorespiration CO_2_ pump further magnified the ammonia imbalance, hence promoting the evolution of C_4_ metabolism, as in the case in Type II (*41*). Although the C_4_ photosynthesis is incomplete and the activities of C_4_ enzymes are low in Type II (*90, 94*), the increased C_4_ cycle might alleviate the decrease in photosynthetic rate caused by the decreased *g_smax_*. In C_4_-like and C_4_ species, a complete and highly active C_4_ cycle (*90*), in particular caused by the increased expression levels and enzymes activities of CA and PEPC, can enable maintenance of high photosynthetic rate even though the reduced intercellular CO_2_ concentration (*2, 4, 17*), which allows *g_smax_* further decreas. This is consistent with the earlier finding that when the limitation on photosynthetic rate by CO_2_ supply becomes less, *g_smax_* dominated by *SD* will reduce (*16*).

### C_4_ evolution select *STOMAGEN* homologs as a molecular trigger to alter *g_smax_*

Stomatal development is controlled by a molecular signaling network composed of many interacting components (*28*). In theory, any gene controlling stomatal development could be potentially selected to alter stomatal patterning and affect g_*smax*_ during the emergence of C_4_ photosynthesis (*15, 80*). In fact, the alterations of stomata patterning under different conditions are indeed caused by different molecular pathways (*24, 95, 96*). Here we show that during C_4_ evolution, the decreased expression of *FSTOMAGEN* (Fig 3c,e, Fig 5b,c,d), a *STOMAGEN*-like gene, is selected to decrease *SD* and *g_smax_* in the *Flaveria* genus and likely in other lineage as well (Fig 6). We used multiple evidences to support this notion. In our experiments, *FSTOMAGEN* strongly and positively regulates stomatal development when either synthetic FSTOMAGEN peptide was applied *in vitro* or *FSTOMAGEN* was over-expressed *in vivo* (Fig 4). Besides *FSTOMAGEN*, we do notice that other genes related to stomatal development were also differentially expressed between C_3_ and C_4_. However, these genes showed either less changes or the pattern of changes did not result in the observed changes in *SD*, as in the case of *EPF1* and *TMM* homologs (Fig 3c,d). For example, *EPF1* homologs negatively regulates stomatal development (*33, 97-99*), but its expression is lower in C_4_, which contradicts with the lower *SD* in C_4_. The decrease in *TMM* and *EPF1* expression levels in C_4_ may be due to the co-regulation and feedback of these stomatal regulators, as previous work shows that overexpression of *STOMAGEN* can increase the expression of other stomata regulatory genes (*59*).

The decreased *FSTOMATGEN* expression might be a major factor for the increased iWUE. The iWUE of C_4_ relative to that of C_3_ increased significantly, partly contributed by a lower *g_s_* (Supplementary fig 1c,d) (*4*). Since the magnitude of the decrease in *g_s_* is similar to the magnitude of the decrease in *g_smax_*, the increased iWUE of C_4_ species can be at least partially attributed to the decreased *g_smax_* (Fig 1; Supplementary fig 1c,d). Because *FSTOMAGEN* induced changes in *SD* and hence *g_smax_*, the decreased *FSTOMAGEN* expression might have contributed to the increased iWUE in C_4_ plants. This is significant for improving iWUE in C_4_ engineering. Many studies suggest that genes related to stomatal development not only affect stomatal development (*100, 101*), but also affect photosynthetic apparatus and the development of photosynthetic tissues (*100*). Therefore, identifying genes that can alter stomata without negatively influencing other parts in C_4_ engineering is required. Results from this study suggests that *FSTOMAGEN* is one such gene, since *FSTOMAGEN* was already selected during the evolution of C_4_ photosynthesis.

## Acknowledgements

We thank Prof. Colin Osborne and Julie Gray for constructive comments on our revised manuscript. The work was funded by Strategic Priority Research Program of the Chinese Academy of Sciences (grant number: XDB27020105) and the general program of National Science Foundation of China (31870214). We thank Prof. Rowan F. Sage (University of Toronto) for providing the *Flaveria* cuttings. We thank Dr. Matt stata for helping cutting of *Flaveria*.

## Author contributions

YYZ and XGZ designed the study. YYZ observed and measured the stomatal characteristic. YYZ carried out the molecular experiment and the treatment of synthetic peptides in vivo. FFM conducted the transgene experiment. GYC guided the treatment of peptides in vivo. Amy MJL analyzed the one kp RNA-seq data. YYZ and XGZ interpreted results and wrote the article.

## Legends for Supplemental Figures

**Supplementary fig 1. a**, Stomatal conductance (*g_s_*) responses to net photosynthesis rate (*A*) under saturated light. The circles on the graph indicate larger *g_s_* values in different photosynthetic types (Data from (*4, 83*)). **b**, The correlation of *g_smax_* (abaxial + adaxial) and *g_s_*. **c**, G_s_, *A and iWUE* under saturated light between C_3_ (rob, pri,cro) and C_4_ (bid, tri and koc) *Flaveria* species. (Data from (*4, 83*)). **d**, G_s_, *A and iWUE* under growth light between C_3_ (rob) and C_4_ (bid) *Flaveria* species (n=6, biologically independent replicates). The asterisks represent statistically significant differences (P<0.05, t-test, two-tailed distribution, unpaired).

**Supplementary fig 2**. *g_smax_* on the adaxial surface of leaves in *Flaveria*. **a**,**b**, The difference in *g_smax_* in species with different photosynthetic types grown outdoors (**a**) or in greenhouse (**b**)**. c**,**d**, The changes of *g_smax_* on the adaxial leaf surface from C_3_ to C_4_ species in the *Flaveria* genus grown outdoors (**c**) and in greenhouse (**d**). (n=3, biologically independent replicates). **e**, Correlation of *g_smax_* and *g_s_* on the adaxial surface of leaves. The asterisks represent statistically significant differences, one-way ANOVA: *, *P* < 0.05, **, *P* < 0.01, ***, *P* < 0.001, ****, *P* < 0.0001; Different letters represent statistically significant differences, Duncan’ s multiple range test (α = 0.05)).

**Supplementary fig 3**. *SD* on the adaxial surface of leaves in *Flaveria*. **a**,**b**, The difference in *SD* in species with different photosynthetic types grown outdoors (**a**) or in greenhouse (**b**)**. c**,**d**, The changes of *SD* on the adaxial leaf surface from C_3_ to C_4_ species in the *Flaveria* genus grown outdoors (**c**) and in greenhouse (**d**). (n=3, biologically independent replicates). **e**, Correlation of *g_smax_* and *SD* on the adaxial surface of leaves. **f**, Correlation of *g_smax_* and *l* on the adaxial surface of leaves. **g**, Correlation of *l* and *SD* on the adaxial surface of leaves. The asterisks represent statistically significant differences, one-way ANOVA: *, *P* < 0.05, **, *P* < 0.01, ***, *P* < 0.001, ****, *P* < 0.0001; Different letters represent statistically significant differences, Duncan’ s multiple range test (α = 0.05)).

**Supplementary fig 4**. *SD* in *rob* (C_3_), *ram* (intermediate) and *bid* (C_4_) grown from seeds. The *SD* on abaxial (ab) (n=5, biologically independent replicates) and adaxial (ad) (n=4, biologically independent replicates) surfaces were observed in *rob*, *ram* and *bid* grown in greenhouse. Different colours represent different photosynthetic types. Different letters represent statistically significant differences (one-way ANOVA, Duncan’s multiple range test (α = 0.05)).

**Supplementary fig 5**. Conservative analysis of amino acid sequences for EPF1 and EPF2. **a**. Comparison of amino acid sequences of homologs of EPF1 between different *Flaveria* species and Arabidopsis. **b**. Comparison of amino acid sequences of homologs of EPF2 between *Flaveria* species and Arabidopsis. White letters with red background represent amino acids conserved across all species. Red letters represent amino acids with similar biochemical property.

**Supplementary fig 6**. Hydrophobicity analysis of proteins. **a**, the protein of stomagen (left) and the protein of FSTOMAGEN (right). **b**, the protein of EPF1 (left) and the protein of FEPF1 (right). **c**, the protein of EPF2 (left) and the protein of FEPF2 (right).

**Supplementary fig 7**. Assessment of the stability of expression levels of reference genes. **a**,**b**, Comparison of the stability of expression levels of candidate reference genes in the *Flaveria* species by RNA-seq. **c**, Gel electrophoresis showing the specificity of the primers for candidate reference genes. From left to right, *ACT7*, *EF1a*, *J3*, *PNdO*, *CPN20*, *UBC9*, *GAPDH*, *F2N1.5*, *PSRP4*, *CACMS*, *HEME2*, and *UBQ11*. **d**, The coefficient of variations (CV) for the candidate reference genes. **e**, The expression stability for candidate reference genes calculated by Genorm. **f**, The expression stability for candidate reference genes calculated by Normfinder.

**Supplementary fig 8**. The expression levels of stomatal development related genes in *Flaveria* genus. Expression levels of stomatal development related genes in species with different photosynthetic types in the *Flaveria* genus by RNA-seq. Data from (*41*).

**Supplementary fig 9**. Relative expression levels of *FSTOMAGEN* and *SD* for *rob* (C_3_) under different light conditions. (A) Relative expression levels for *FSTOMAGEN* under two different light conditions. (B) *SD* under two different light conditions. Natural light: the maximal photosynthetic photon flux density in the greenhouse was about 1300~1400 mmol m^−2^ s^−1^. Light in the phytotron: 20~100 mmol m^−2^ s^−1^. Reference gene used to calculate the expression level was *ACT7*. The asterisks represent statistically significant differences (*P*<0.05, t-test, two-tailed distribution, unpaired).

**Supplementary fig 10**. Relative expression levels of stomata development related genes determined with different reference genes and different primers. **a, b, c**, Quantification with different primers and the same *EF1a* reference gene. Relative expressions shown in **a and b** were determined using the same primer pairs (Frob_EF1a, Fram_EF1a, Fbid_EF1a) while that in **c** used other primer pairs (EF1a) (See details of the primers used in Table S2). **d**, Relative expression levels determined with the gene *J3* used as the reference gene.

**Supplementary fig 11**. Transcript abundance of genes in the C_4_ cycle and C_4_ related transporters in *Flaveria* genus. Data was from the One Thousand Plants (1KP)(*42, 65*). J represents juvenile leaves and M represents mature leaves. Different colours represent species with different photosynthetic types in the *Flaveria* genus. Red colour represents C_3_ species, grey colour represents C_3_-C_4_ species, green colour represents C_4_-like species, and blue colour represents C_4_ species.

**Supplementary fig 12**. Alignment of the functional domain of *STOMAGEN* with transcripts from different *Flaveria* species by Blast. From the top to bottom, *F. robusta*, *F. pringlei*, *F. angustifolia*, *F. ramosissima*, *F. vaginata*, *F. australasica*, *F. bidentis*.

**Supplementary fig 13**. The expression level of *FSTOMAGEN* in different *Flaveria* species based on RT-qPCR (**a**) or RNA-seq (**b**). For RT-qPCR, *pri* (n=5), *rob* (n=13), *ang* (n=3), *ram* (n=5), *vag* (n=3), *tri* (n=4), *aus* (n=3), *bid* (n=4) and *koc* (n=3). For RNA-seq, all species n=4. The relative expression indicates the measurements with RT-qPCR. RPKM indicates RNA-seq, data from (*41*). **c**, The relationship between *SD* and *FSTOMAGEN* expression in *Flaveria* species with different photosynthetic types.

**Supplementary fig 14**. Comparison of the expression level of *FEPF1* and *FEPF2* between *rob* and *bid Flaveria* species by RT-qPCR or RNA-seq. RNA-seq data from (*41*). *FEPF1* and *FEPF2* indicates the homologs of *EPF1* and *EPF2*.

**Supplementary fig 15**. The analysis for the *SD* of T2 generation of transgenic Arabidopsis (AFSTO and FSTO). **a**, Comparison of *SD* between WT and transgenic Arabidopsis AR, FSTO respectively. **b**, RNA level identification of transgenic Arabidospsis. Two independent lines was analyzed for each trangene event.

**Supplementary fig 16**. The increase of *SD* in cotyledon of *F. bidentis* (*bid*) with in vitro application of FSTOMAGEN. Photographs of the leaf surface in the untreated (control) and treated bid plants. scale bar, 20 mm.

**Supplementary fig 17.a**, Photographs of leaf surface taken with differential interference contrast microscope showing the increased stomata density in F. bidentis (bid) leaves with FSTOMAGEN applied. Scale bar represents 20 μm. **b**, Photographs of leaf surface taken with differential interference contrast microscope showing the increased stomata density in Arabidopsis leaves with FSTOMAGEN applied. Scale bar represents 20 μm.

**Supplementary fig 18**. Phylogenetic tree for FSTOMAGEN. blue represents dicotyledon, red represents monocotyledon and pink represents the *Flaveria* species. The gene tree of STOMAGEN based on bayesian-inference method.

**Supplementary fig 19. a**, Schematic of the progresssion of C_4_ evolution in *Flaveria* genus. **b**, A stomatal graph taken with a light microscope showing stomatal parameters for *Flaveria* species which are needed to be measured to calculate the *g_smax_*. The graph of stomata is *rob*. *l* (stomatal length), PL (pore length) and GCW (guard cell width). **c**, Correlation of the guard cell width and stomatal length (left), correlation of the stomatal length and pore length (right).

**Supplementary fig 20**. Photographs showing clustered stomata on a leaf for the transgenic Arabidopsis. Scar bar represents 10 um.

**Supplementary table 1**. Statistics for the mapping of the RNA-seq data of the juvenile and mature leaves in different species of the *Flaveria* genus.

**Supplementary table 2**. Primers used in this study

**Supplementary table 3**. The effect and accession number of genes related to stomatal development in Arabidopsis and rice and maize.

**Supplementary table 4**. The N50 of the mapping of the RNA-SEQ data taken from 1KP database for the juvenile and mature leaves

**Supplementary table 5**. The mapping rates of the mapping of the RNA-SEQ data taken from 1KP database for the juvenile and mature leaves.

## Reference

1. F. I. W. Alistair M. Hetherington, The role of stomata in sensing and driving environmental change.pdf. Nature 424, 901–908 (2003).

2. S. H. H. Taylor, S. P. Rees, M Ripley, B. S. Woodward, F. I. Osborne, C. P., Ecophysiological traits in C_3_ and C_4_ grasses: a phylogenetically controlled screening experiment. New Phytol 185, 780–791 (2010).

3. C. P. Osborne, L. Sack, Evolution of C_4_ plants: a new hypothesis for an interaction of CO_2_ and water relations mediated by plant hydraulics. Philos Trans R Soc Lond B Biol Sci 367, 583–600 (2012).

4. P. J. Vogan, R. F. Sage, Water-use efficiency and nitrogen-use efficiency of C_3_-C_4_ intermediate species of Flaveria Juss. (Asteraceae). Plant, cell & environment 34, 1415–1430 (2011).

5. H. Zhou, B. R. Helliker, M. Huber, A. Dicks, E. Akcay, C_4_ photosynthesis and climate through the lens of optimality. Proc Natl Acad Sci U S A 115, 12057–12062 (2018).

6. G. Reeves et al., Natural Variation within a Species for Traits Underpinning C_4_ Photosynthesis. Plant physiology 177, 504–512 (2018).

7. B. P. Williams, I. G. Johnston, S. Covshoff, J. M. Hibberd, Phenotypic landscape inference reveals multiple evolutionary paths to C_4_ photosynthesis. elife 2, e00961 (2013).

8. S. H. Taylor, B. S. Ripley, F. I. Woodward, C. P. Osborne, Drought limitation of photosynthesis differs between C(3)and C(4)grass species in a comparative experiment. Plant, cell & environment 34, 65–75 (2011).

9. F. Kocacinar, Photosynthetic, hydraulic and biomass properties in closely related C_3_ and C_4_species. Physiologia Plantarum 153, 454–466 (2015).

10. B. Uzilday, I. Turkan, R. Ozgur, A. H. Sekmen, Strategies of ROS regulation and antioxidant defense during transition from C(3) to C(4) photosynthesis in the genus Flaveria under PEG-induced osmotic stress. J Plant Physiol 171, 65–75 (2014).

11. S. A. Aubry, O. Reyna-Llorens, I. Smith-Unna, R. D. Hibberd, J. M. Genty, B., A Specific Transcriptome Signature for Guard Cells from the C_4_ Plant Gynandropsis gynandra. Plant physiology 170, 1345–1357 (2016).

12. R. F. Sage, The evolution of C_4_ photosynthesis. New Phytologist 161, 341–270 (2004).

13. A. M. Powell, Systematics of Flaveria (Flaveriinae--Asteraceae). Annals of the Missouri Botanical Garden 65, 590–636 (178).

14. P. J. Franks, P. L. Drake, D. J. Beerling, Plasticity in maximum stomatal conductance constrained by negative correlation between stomatal size and density: an analysis using Eucalyptus globulus. Plant, cell & environment 32, 1737–1748 (2009).

15. T. Doheny-Adams, L. Hunt, P. J. Franks, D. J. Beerling, J. E. Gray, Genetic manipulation of stomatal density influences stomatal size, plant growth and tolerance to restricted water supply across a growth carbon dioxide gradient. Philos Trans R Soc Lond B Biol Sci 367, 547–555 (2012).

16. P. J. F. a. D. J. Beerling, Maximum leaf conductance driven by CO_2_ effects on stomatal size and density over geologic time. PNAS 106, 10343–10347 (2009).

17. S. H. Taylor et al., Photosynthetic pathway and ecological adaptation explain stomatal trait diversity amongst grasses. New Phytol 193, 387–396 (2012).

18. W. M. Laetsch, The C_4_’ syndrome: a structural analysis. Ann. Rev. Plant Physiol. 25, 27–52 (1974).

19. D. A. Way, What lies between : the evolution of stomatal traits on the road to C_4_ photosynthesis. New phytologist 193, 291–293 (2012).

20. T. W. Ocheltree, J. B. Nippert, P. V. Prasad, Changes in stomatal conductance along grass blades reflect changes in leaf structure. Plant, cell & environment 35, 1040–1049 (2012).

21. V. S. R. D. a. M. Santakumari, Stomatal characteristics of some dicotyledonous plants in relation to the C_4_ and C_3_ pathways of photosynthesis. Plant and cell physiology 18, 935–938 (1977).

22. E. J. Edwards, C. J. Still, Climate, phylogeny and the ecological distribution of C_4_ grasses. Ecol Lett 11, 266–276 (2008).

23. E. J. Edwards, C. J. Still, M. J. Donoghue, The relevance of phylogeny to studies of global change. Trends Ecol Evol 22, 243–249 (2007).

24. E. T. Hamanishi, B. R. Thomas, M. M. Campbell, Drought induces alterations in the stomatal development program in Populus. J Exp Bot 63, 4959–4971 (2012).

25. Q. Cai, C. Ji, Z. Yan, X. Jiang, J. Fang, Anatomical responses of leaf and stem of Arabidopsis thaliana to nitrogen and phosphorus addition. J Plant Res 130, 1035–1045 (2017).

26. Z. Xu, G. Zhou, Responses of leaf stomatal density to water status and its relationship with photosynthesis in a grass. J Exp Bot 59, 3317–3325 (2008).

27. N. Sekiya, K. Yano, Stomatal density of cowpea correlates with carbon isotope discrimination in different phosphorus, water and CO_2_ environments. New Phytol 179, 799–807 (2008).

28. N. Zoulias, E. L. Harrison, S. A. Casson, J. E. Gray, Molecular control of stomatal development. Biochem J 475, 441–454 (2018).

29. G. Lin et al., A receptor-like protein acts as a specificity switch for the regulation of stomatal development. Genes Dev 31, 927–938 (2017).

30. E. D. Shpak, J. M. McAbee, L. J. Pillitteri, K. U. Torii, Stomatal patterning and differentiation by synergistic interactions of receptor kinases. Science 309, 290–293 (2005).

31. J. S. Lee et al., Direct interaction of ligand-receptor pairs specifying stomatal patterning. Genes Dev 26, 126–136 (2012).

32. T. Shimada, S. S. Sugano, I. Hara-Nishimura, Positive and negative peptide signals control stomatal density. Cell Mol Life Sci 68, 2081–2088 (2011).

33. K. Hara, R. Kajita, K. U. Torii, D. C. Bergmann, T. Kakimoto, The secretory peptide gene EPF1 enforces the stomatal one-cell-spacing rule. Genes Dev 21, 1720–1725 (2007).

34. L. Hunt, J. E. Gray, The signaling peptide EPF2 controls asymmetric cell divisions during stomatal development. Curr Biol 19, 864–869 (2009).

35. L. J. Pillitteri, K. U. Torii, Breaking the silence: three bHLH proteins direct cell-fate decisions during stomatal development. Bioessays 29, 861–870 (2007).

36. S. S. Sugano et al., Stomagen positively regulates stomatal density in Arabidopsis. Nature 463, 241–244 (2010).

37. T. Kondo et al., Stomatal density is controlled by a mesophyll-derived signaling molecule. Plant Cell Physiol 51, 1–8 (2010).

38. L. Hunt, K. J. Bailey, J. E. Gray, The signalling peptide EPFL9 is a positive regulator of stomatal development. New Phytol 186, 609–614 (2010).

39. U. Schluter, A. P. Weber, The Road to C_4_ Photosynthesis: Evolution of a Complex Trait via Intermediary States. Plant Cell Physiol 57, 881–889 (2016).

40. D. Heckmann et al., Predicting C_4_ photosynthesis evolution: modular, individually adaptive steps on a Mount Fuji fitness landscape. Cell 153, 1579–1588 (2013).

41. J. Mallmann et al., The role of photorespiration during the evolution of C_4_ photosynthesis in the genus Flaveria. Elife 3, e02478 (2014).

42. M. J. Lyu et al., RNA-Seq based phylogeny recapitulates previous phylogeny of the genus Flaveria (Asteraceae) with some modifications. BMC Evol Biol 15, 116 (2015).

43. A. D. M. A. N. G. Dengler, Key innovations in the evolution of Kranz anatomy and C_4_ vein pattern in Flaveria (Asteraceae). American Journal of Botany 94, 382–399 (2007).

44. S. Schulze et al., Evolution of C_4_ photosynthesis in the genus flaveria: establishment of a photorespiratory CO_2_ pump. Plant Cell 25, 2522–2535 (2013).

45. M. Stata et al., Mesophyll Chloroplast Investment in C_3_, C_4_ and C2 Species of the Genus Flaveria. Plant Cell Physiol 57, 904–918 (2016).

46. P. A. Christin, C. P. Osborne, R. F. Sage, M. Arakaki, E. J. Edwards, C(4) eudicots are not younger than C(4) monocots. J Exp Bot 62, 3171–3181 (2011).

47. U. Gowik, A. Bräutigam, K. L. Weber, A. P. M. Weber, P. Westhoff, Evolution of C_4_ Photosynthesis in the Genus Flaveria: How Many and Which Genes Does It Take to Make C_4_? The Plant Cell 23, 2087–2105 (2011).

48. A. D. Mckown, PHYLOGENY OF FLAVERIA (ASTERACEAE) AND INFERENCE OF C_4_ PHOTOSYNTHESIS EVOLUTION. American Journal of Botany 92, 1911–1928 (2005).

49. E. A. Sudderth, R. M. Muhaidat, A. D. McKown, F. Kocacinar, R. F. Sage, Leaf anatomy, gas exchange and photosynthetic enzyme activity in Flaveria kochiana. Funct Plant Biol 34, 118–129 (2007).

50. E. A. Sudderth, F. J. Espinosa-García, N. M. Holbrook, Geographic distributions and physiological characteristics of co-existing Flaveria species in south-central Mexico. Flora - Morphology, Distribution, Functional Ecology of Plants 204, 89–98 (2009).

51. F. Kocacinar, A. D. McKown, T. L. Sage, R. F. Sage, Photosynthetic pathway influences xylem structure and function in Flaveria (Asteraceae). Plant, cell & environment 31, 1363–1376 (2008).

52. N. Nakamura, M. Iwano, M. Havaux, A. Yokota, Y. N. Munekage, Promotion of cyclic electron transport around photosystem I during the evolution of NADP-malic enzyme-type C_4_ photosynthesis in the genus Flaveria. New Phytol 199, 832–842 (2013).

53. R. F. Sage, A portrait of the C_4_ photosynthetic family on the 50th anniversary of its discovery: species number, evolutionary lineages, and Hall of Fame. J Exp Bot 68, 4039–4056 (2017).

54. Y. Y. Taniguchi et al., Dynamic changes of genome sizes and gradual gain of cell-specific distribution of C_4_ enzymes during C_4_ evolution in genus Flaveria. Plant Genome, e20095 (2021).

55. R. F. Sage, R. Khoshravesh, T. L. Sage, From proto-Kranz to C_4_ Kranz: building the bridge to C_4_ photosynthesis. J Exp Bot 65, 3341–3356 (2014).

56. S. R. Christopher M. Hylton, Alison M. Smith, D. Alan Jones and Harold W. Woolhouse, Glycine decarboxylase is confined to the bundle-sheath cells of leaves of C_3_-C_4_ intermediate species. planta 175, 452–459 (1988).

57. L. J. Pillitteri, K. U. Torii, Mechanisms of stomatal development. Annu Rev Plant Biol 63, 591–614 (2012).

58. S. Ohki, M. Takeuchi, M. Mori, The NMR structure of stomagen reveals the basis of stomatal density regulation by plant peptide hormones. Nat Commun 2, 512 (2011).

59. M. Hronkova et al., Light-induced STOMAGEN-mediated stomatal development in Arabidopsis leaves. J Exp Bot 66, 4621–4630 (2015).

60. J. S. Lee et al., Competitive binding of antagonistic peptides fine-tunes stomatal patterning. Nature 522, 439–443 (2015).

61. J. Lu et al., Homologous genes of epidermal patterning factor regulate stomatal development in rice. J Plant Physiol 234-235, 18–27 (2019).

62. L. Xuxin, The Effects for Stomata Developmental on Arabidopsis by Overexpression of Rice EPFL Family Genes. Molecular Plant Breeding 15, 2042–2047 (2017).

63. X. Yin et al., CRISPR-Cas9 and CRISPR-Cpf1 mediated targeting of a stomatal developmental gene EPFL9 in rice. Plant Cell Rep 36, 745–757 (2017).

64. J. Y. Zhang, S. B. He, L. Li, H. Q. Yang, Auxin inhibits stomatal development through MONOPTEROS repression of a mobile peptide gene STOMAGEN in mesophyll. Proc Natl Acad Sci U S A 111, E3015–3023 (2014).

65. M. A. Lyu et al., The coordination of major events in C_4_ photosynthesis evolution in the genus Flaveria. Sci Rep 11, 15618 (2021).

66. K. Billakurthi et al., Transcriptome dynamics in developing leaves from C_3_ and C_4_ Flaveria species reveal determinants of Kranz anatomy. bioRxiv, (2018).

67. M. G. Grabherr et al., Full-length transcriptome assembly from RNA-Seq data without a reference genome. Nat Biotechnol 29, 644–652 (2011).

68. B. Li, C. N. Dewey, RSEM: accurate transcript quantification from RNA-Seq data with or without a reference genome. BMC Bioinformatics 12, 323 (2011).

69. L. Wang et al., Comparative analyses of C_4_ and C_3_ photosynthesis in developing leaves of maize and rice. Nat Biotechnol 32, 1158–1165 (2014).

70. X. Robert, P. Gouet, Deciphering key features in protein structures with the new ENDscript server. Nucleic Acids Res 42, W320–324 (2014).

71. D. R. F. Kyte J, A simple method for displaying the hydropathic character of a protein. Journal of Molecular Biology 157, 105–132 (1982).

72. S. Perveen et al., Overexpression of maize transcription factor mEmBP-1 increases photosynthesis, biomass, and yield in rice. J Exp Bot 71, 4944–4957 (2020).

73. D. M. Emms, S. Kelly, OrthoFinder: solving fundamental biases in whole genome comparisons dramatically improves orthogroup inference accuracy. Genome Biol 16, 157 (2015).

74. R. C. Edgar, MUSCLE: multiple sequence alignment with high accuracy and high throughput. Nucleic Acids Res 32, 1792–1797 (2004).

75. A. Stamatakis, RAxML-VI-HPC: maximum likelihood-based phylogenetic analyses with thousands of taxa and mixed models. Bioinformatics 22, 2688–2690 (2006).

76. F. Ronquist et al., MrBayes 3.2: efficient Bayesian phylogenetic inference and model choice across a large model space. Syst Biol 61, 539–542 (2012).

77. S. A. Shiryev, J. S. Papadopoulos, A. A. Schaffer, R. Agarwala, Improved BLAST searches using longer words for protein seeding. Bioinformatics 23, 2949–2951 (2007).

78. X. J. Min, G. Butler, R. Storms, A. Tsang, OrfPredictor: predicting protein-coding regions in EST-derived sequences. Nucleic Acids Res 33, W677–680 (2005).

79. A. L. Ming-Ju et al., The Coordination and Jumps along C_4_ Photosynthesis Evolution in the Genus Flaveria. bioRxiv, 460287 (2018).

80. P. J. Franks, W. Doheny-Adams T, Z. J. Britton-Harper, J. E. Gray, Increasing water-use efficiency directly through genetic manipulation of stomatal density. New Phytol 207, 188–195 (2015).

81. J. L. J. a. T. F. Ø. Claus Lindbjerg Andersen, Normalization of real-time quantitative reverse transcription-PCR data a model-based variance estimation approach to identify genes suited for normalization, applied to bladder and colon cancer data sets. Cancer Res 64, 5245–5250 (2004).

82. D. P. K. Vandesompele J, Pattyn F, Poppe B, Van Roy N, De Paepe A, Speleman F., Accurate normalization of real-time quantitative RT-PCR data by geometric averaging of multiple internal control genes. Genome Biology 3, research0034.0031–0034.0011 (2002).

83. D. A. Way, G. G. Katul, S. Manzoni, G. Vico, Increasing water use efficiency along the C_3_ to C_4_ evolutionary pathway: a stomatal optimization perspective. J Exp Bot 65, 3683–3693 (2014).

84. A. V. a. D. C. Bergmann, Mechanisms of stomatal development: an evolutionary view. EvoDevo 3, (2012).

85. P. J. Franks, D. J. Beerling, CO(2)-forced evolution of plant gas exchange capacity and water-use efficiency over the Phanerozoic. Geobiology 7, 227–236 (2009).

86. I. R. C. G. D. F. S. C. Wong, Stomatal conductance correlates with photosynthetic capacity. Nature 282, 424–426 (1979).

87. Y. Tanaka, S. S. Sugano, T. Shimada, I. Hara-Nishimura, Enhancement of leaf photosynthetic capacity through increased stomatal density in Arabidopsis. New Phytol 198, 757–764 (2013).

88. J. M. Beaulieu, I. J. Leitch, S. Patel, A. Pendharkar, C. A. Knight, Genome size is a strong predictor of cell size and stomatal density in angiosperms. New Phytol 179, 975–986 (2008).

89. P. J. Vogan, M. W. Frohlich, R. F. Sage, The functional significance of C_3_-C_4_ intermediate traits in Heliotropium L. (Boraginaceae): gas exchange perspectives. Plant, cell & environment 30, 1337–1345 (2007).

90. J. W. Maurice S. B. Ku, Ziyu Dai, Rick A. Scott, Chun Chu, and Gerald E. Edwards, Photosynthetic and Photorespiratory Characteristics of Flaveria species. plant physiol 96, 518–528 (1991).

91. O. Keerberg, T. Parnik, H. Ivanova, B. Bassuner, H. Bauwe, C2 photosynthesis generates about 3-fold elevated leaf CO_2_ levels in the C_3_-C_4_ intermediate species Flaveria pubescens. J Exp Bot 65, 3649–3656 (2014).

92. R. F. Sage, T. L. Sage, F. Kocacinar, Photorespiration and the evolution of C_4_ photosynthesis. Annu Rev Plant Biol 63, 19–47 (2012).

93. P. J. Vogan, R. F. Sage, Effects of low atmospheric CO_2_ and elevated temperature during growth on the gas exchange responses of C_3_, C_3_-C_4_ intermediate, and C_4_ species from three evolutionary lineages of C_4_ photosynthesis. Oecologia 169, 341–352 (2012).

94. R. K. M. Maurice S. B. Ku, Robert. Littlejohn, JR., Hitoshi Nakamoto, Donald B. Fisher, And Gerald E. Edwards, Photosynthetic Characteristics of C_3_-C_4_ Intermediate Flaveria Species. Plant Physiol. 71, 944–948 (1983).

95. C. B. Engineer et al., Carbonic anhydrases, EPF2 and a novel protease mediate CO_2_ control of stomatal development. Nature 513, 246–250 (2014).

96. Y. Tanaka, T. Nose, Y. Jikumaru, Y. Kamiya, ABA inhibits entry into stomatal-lineage development in Arabidopsis leaves. Plant J 74, 448–457 (2013).

97. C. Wang et al., PdEPF1 regulates water-use efficiency and drought tolerance by modulating stomatal density in poplar. Plant Biotechnol J 14, 849–860 (2016).

98. R. S. Caine et al., Rice with reduced stomatal density conserves water and has improved drought tolerance under future climate conditions. New Phytol, (2018).

99. J. Hughes et al., Reducing Stomatal Density in Barley Improves Drought Tolerance without Impacting on Yield. Plant physiology 174, 776–787 (2017).

100. G. J. Dow, J. A. Berry, D. C. Bergmann, Disruption of stomatal lineage signaling or transcriptional regulators has differential effects on mesophyll development, but maintains coordination of gas exchange. New Phytol 216, 69–75 (2017).

101. H. Cui et al., SPINDLY, ERECTA, and its ligand STOMAGEN have a role in redox-mediated cortex proliferation in the Arabidopsis root. Mol Plant 7, 1727–1739 (2014).

102. M. Stata, T. L. Sage, R. F. Sage, Mind the gap: the evolutionary engagement of the C_4_ metabolic cycle in support of net carbon assimilation. Curr Opin Plant Biol 49, 27–34 (2019).

