## Supplementary figures and images for "The evolution of stomatal traits along the trajectory towards C_4_ photosynthesis"

### supplementary fig 1

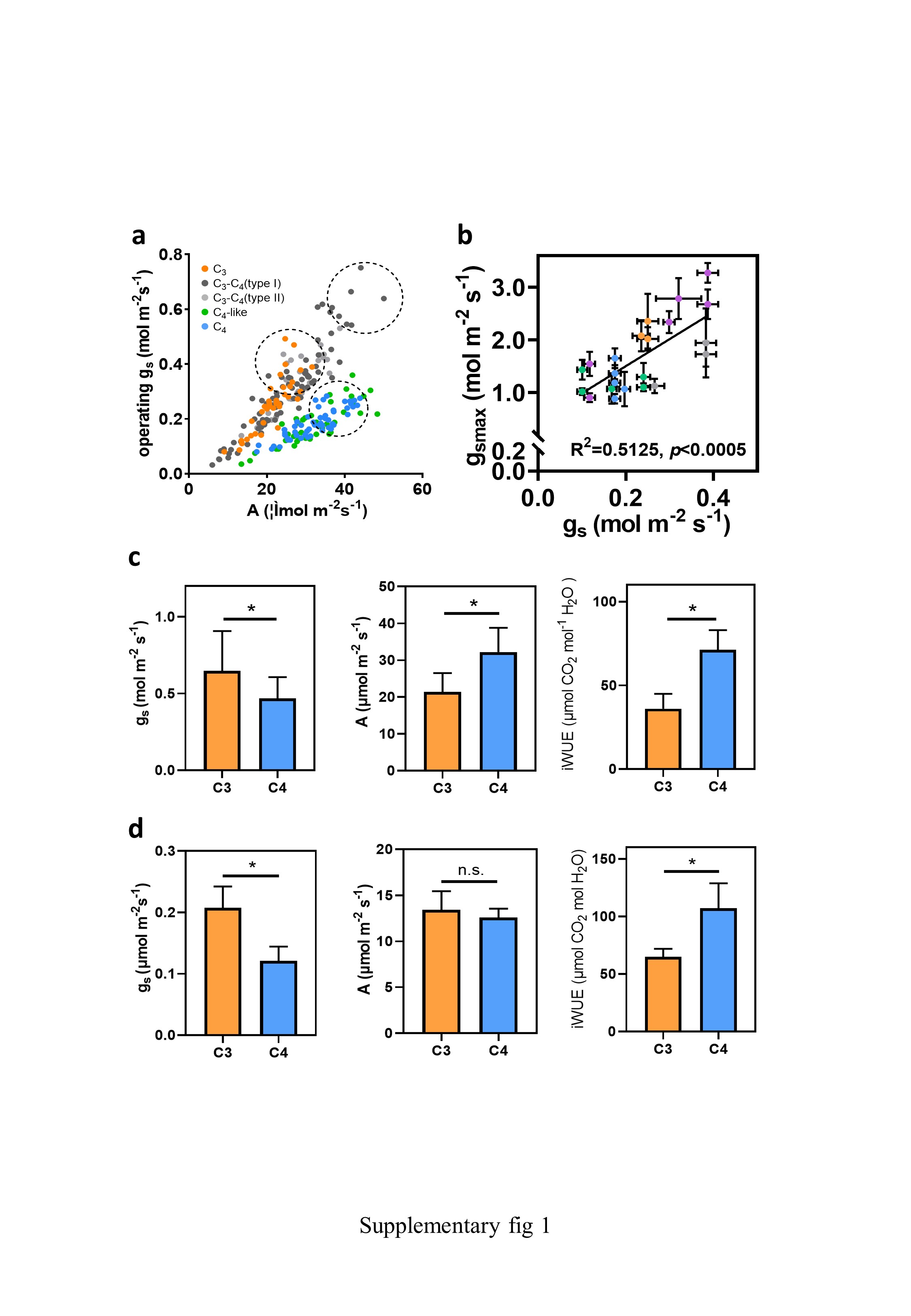

### supplementary fig 2

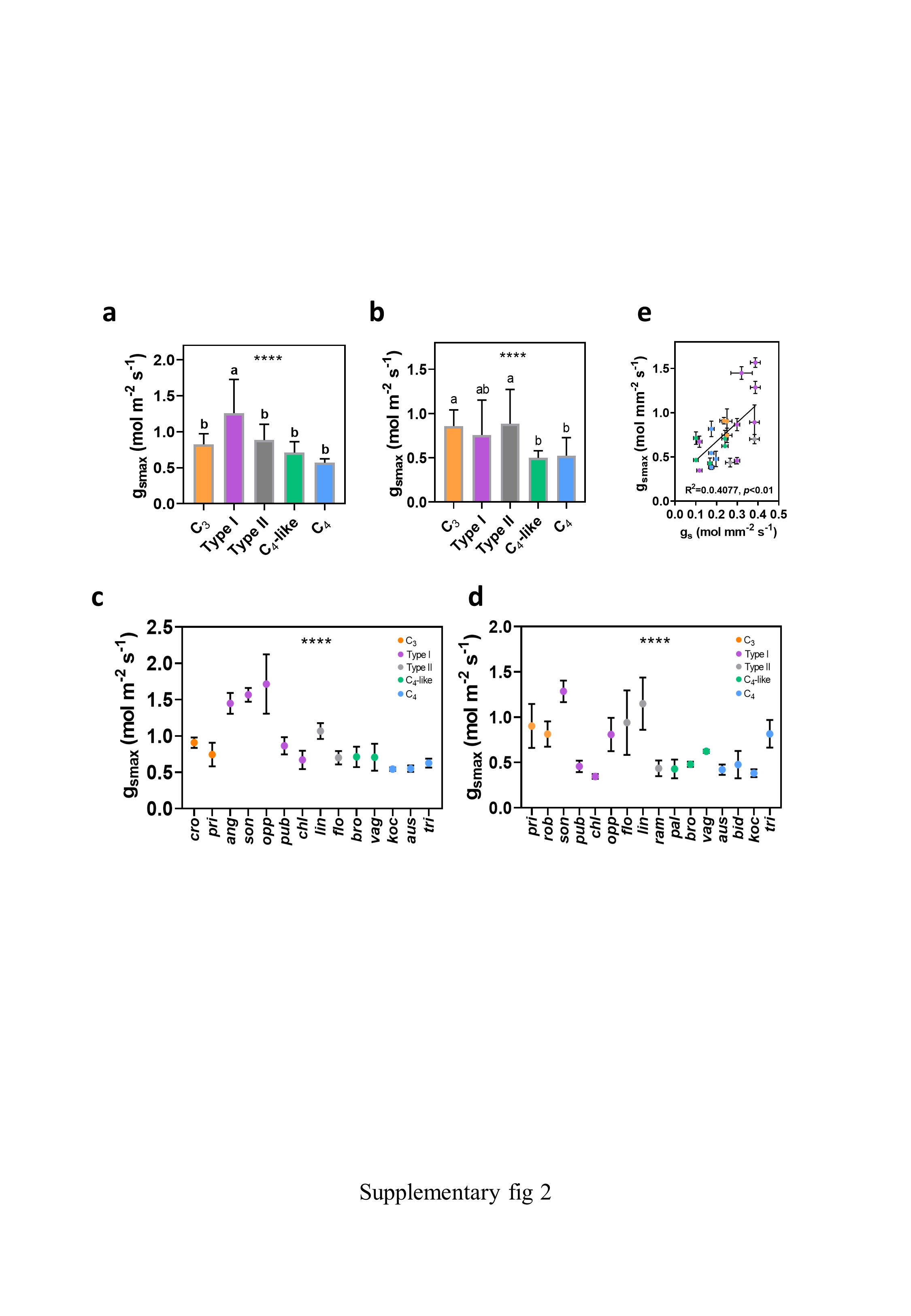

### supplementary fig 3

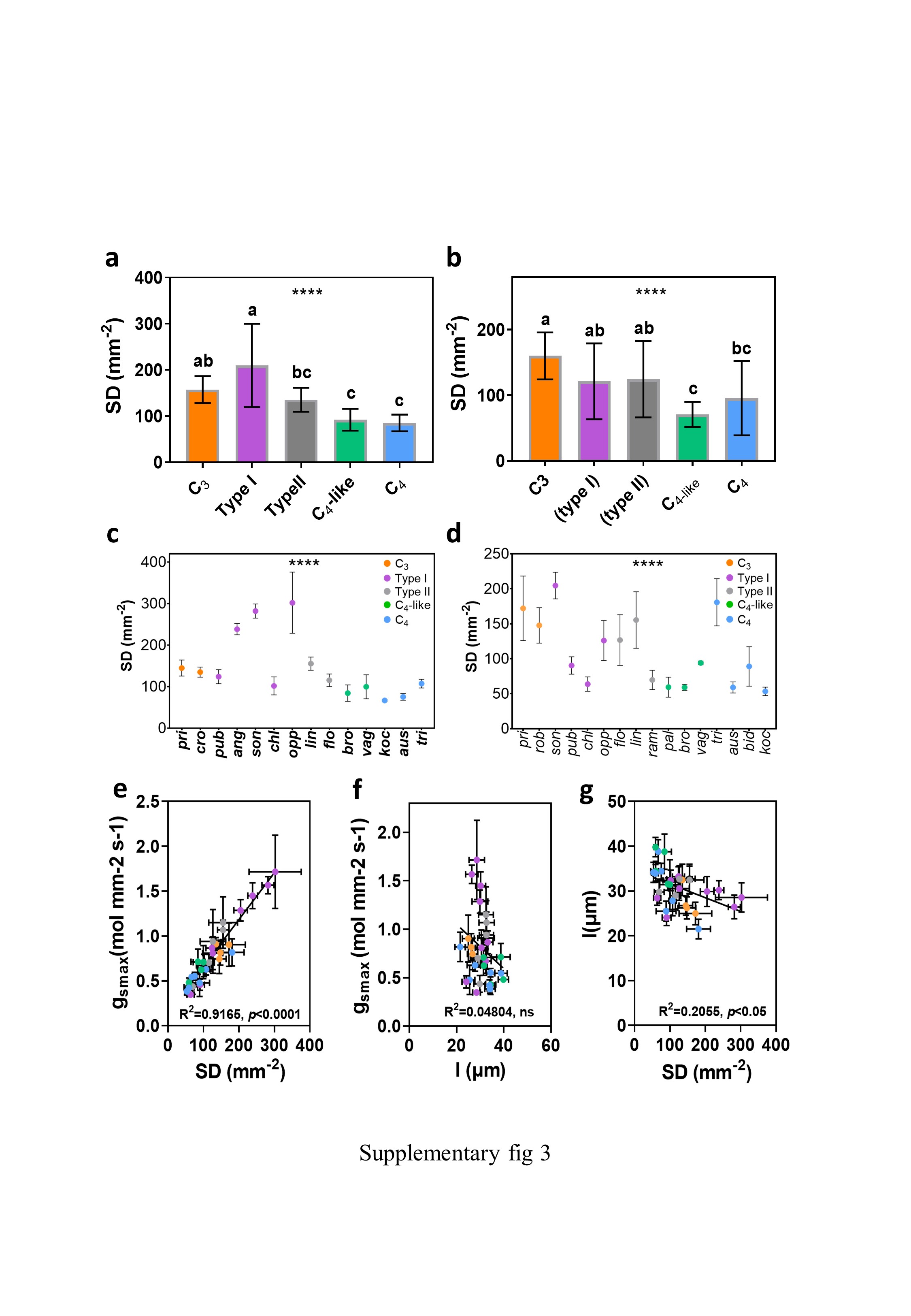

### supplementary fig 4

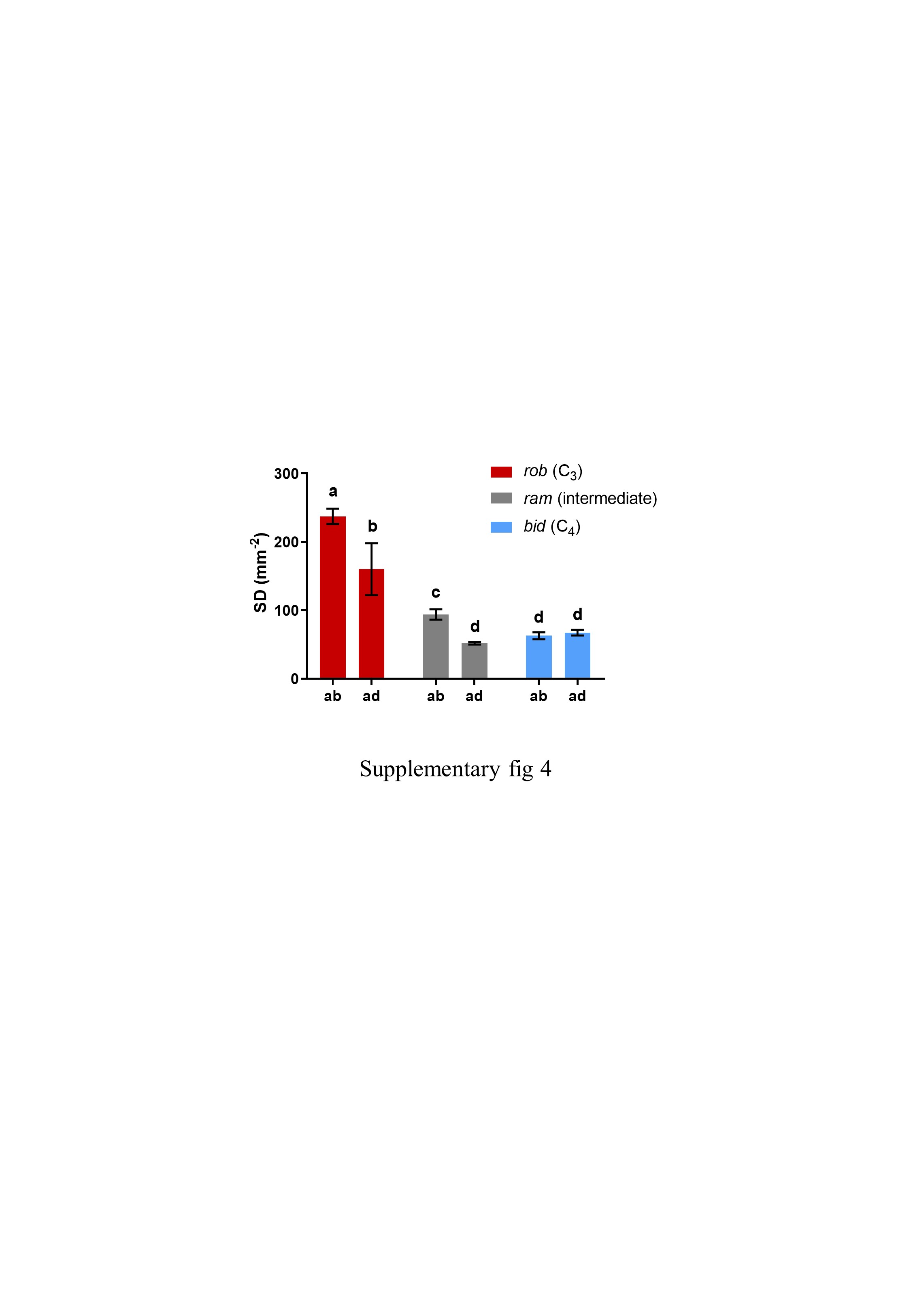

### supplementary fig 5

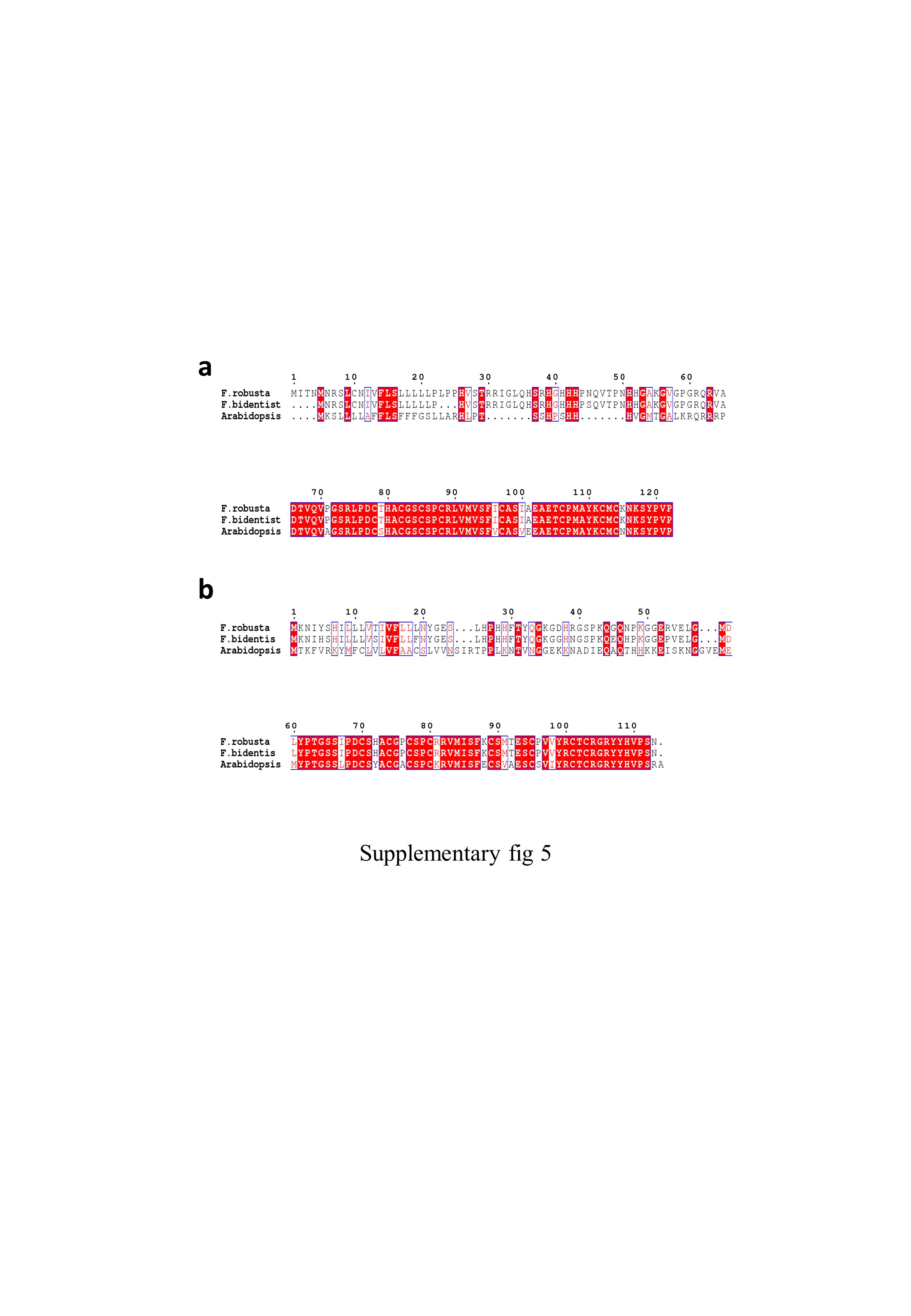

### supplementary fig 6

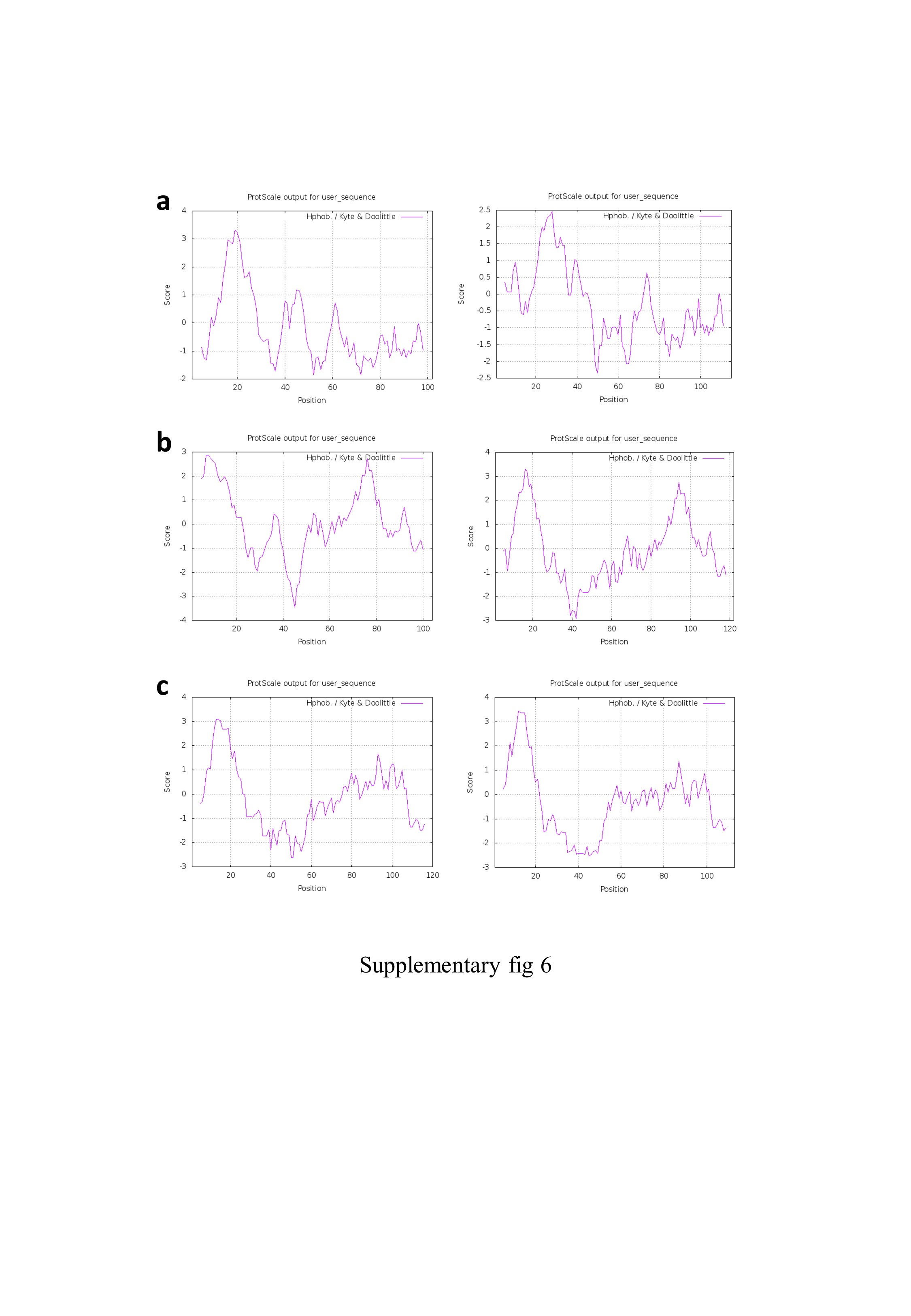

### supplementary fig 7

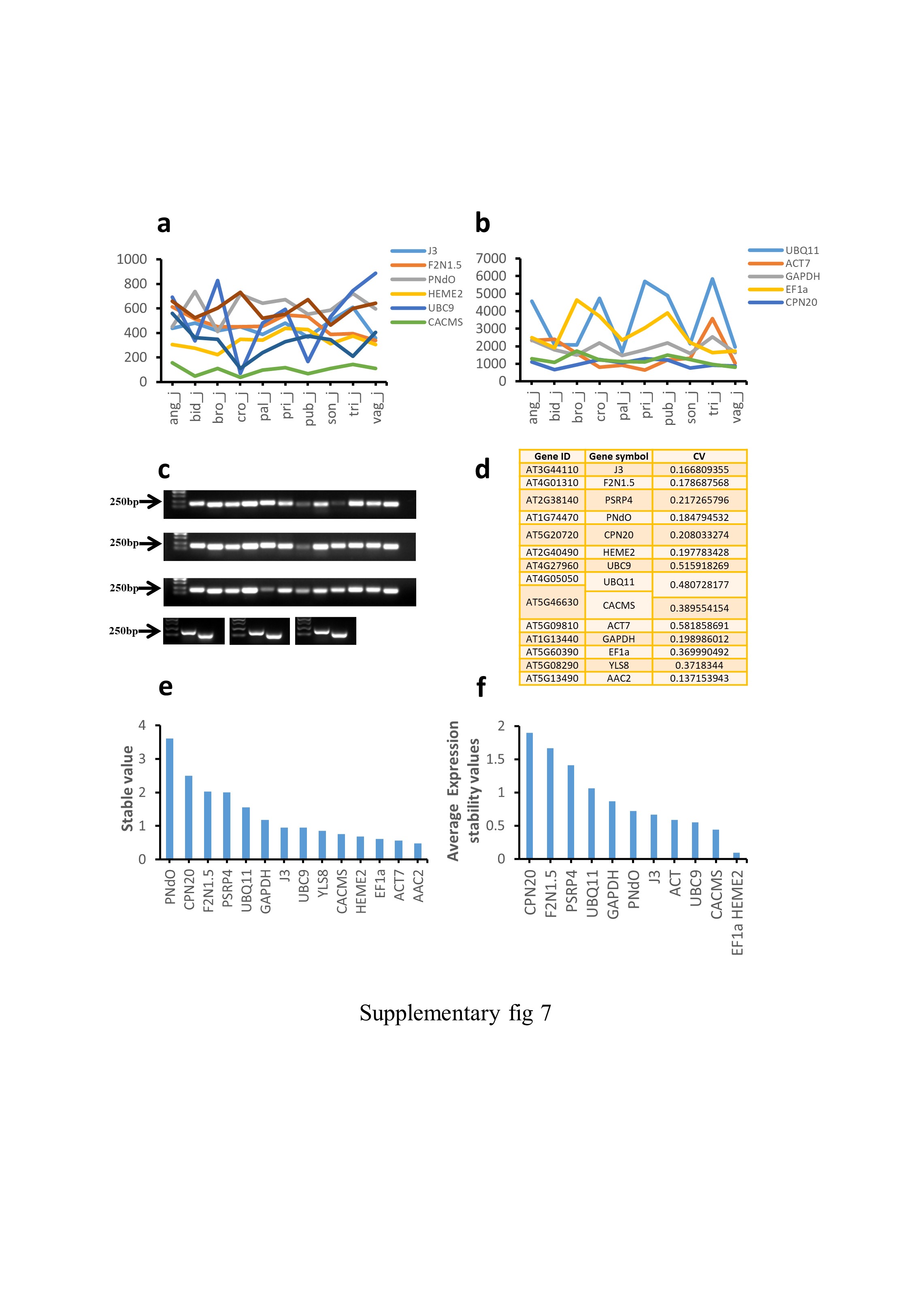

### supplementary fig 8

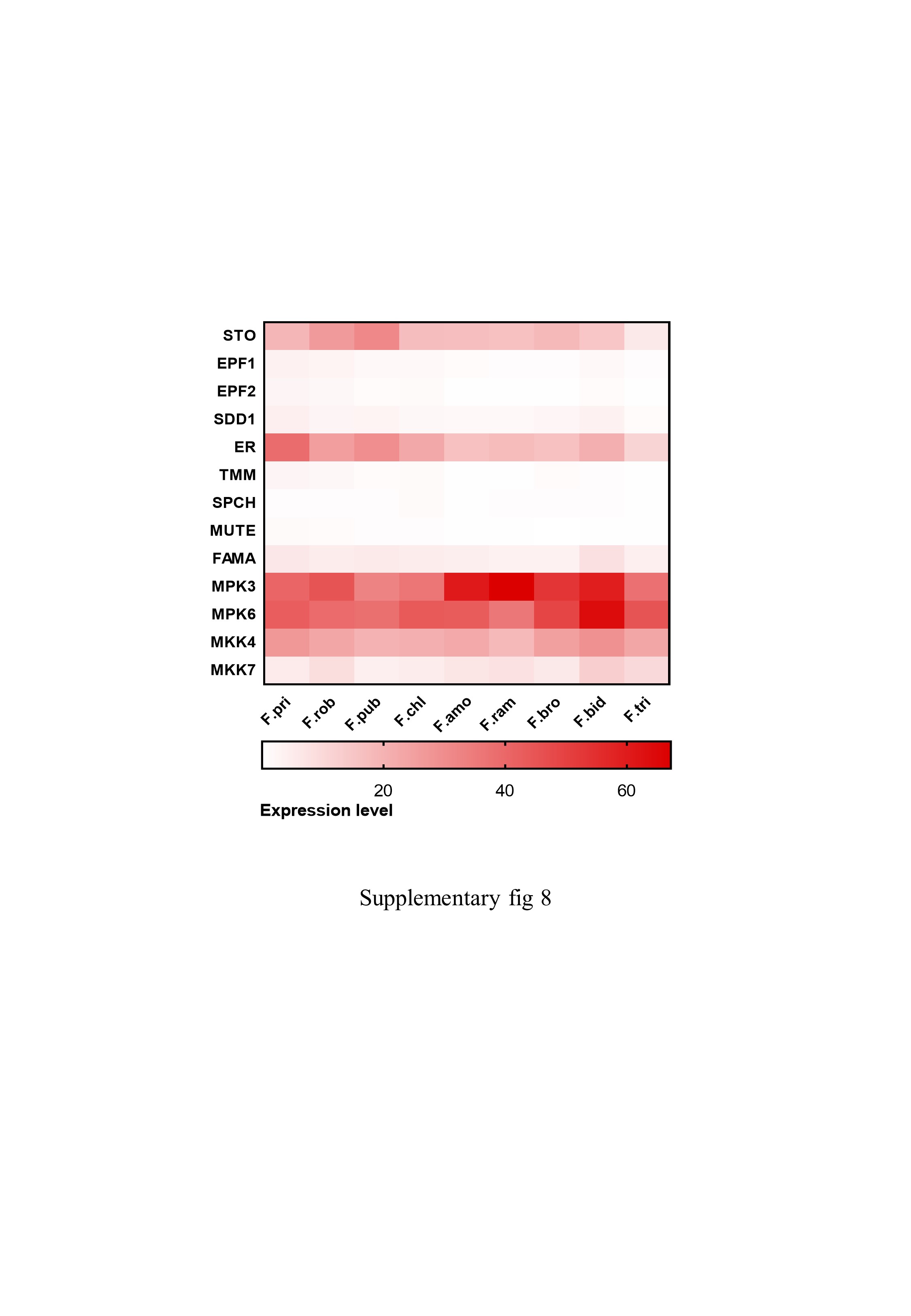

### supplementary fig 9

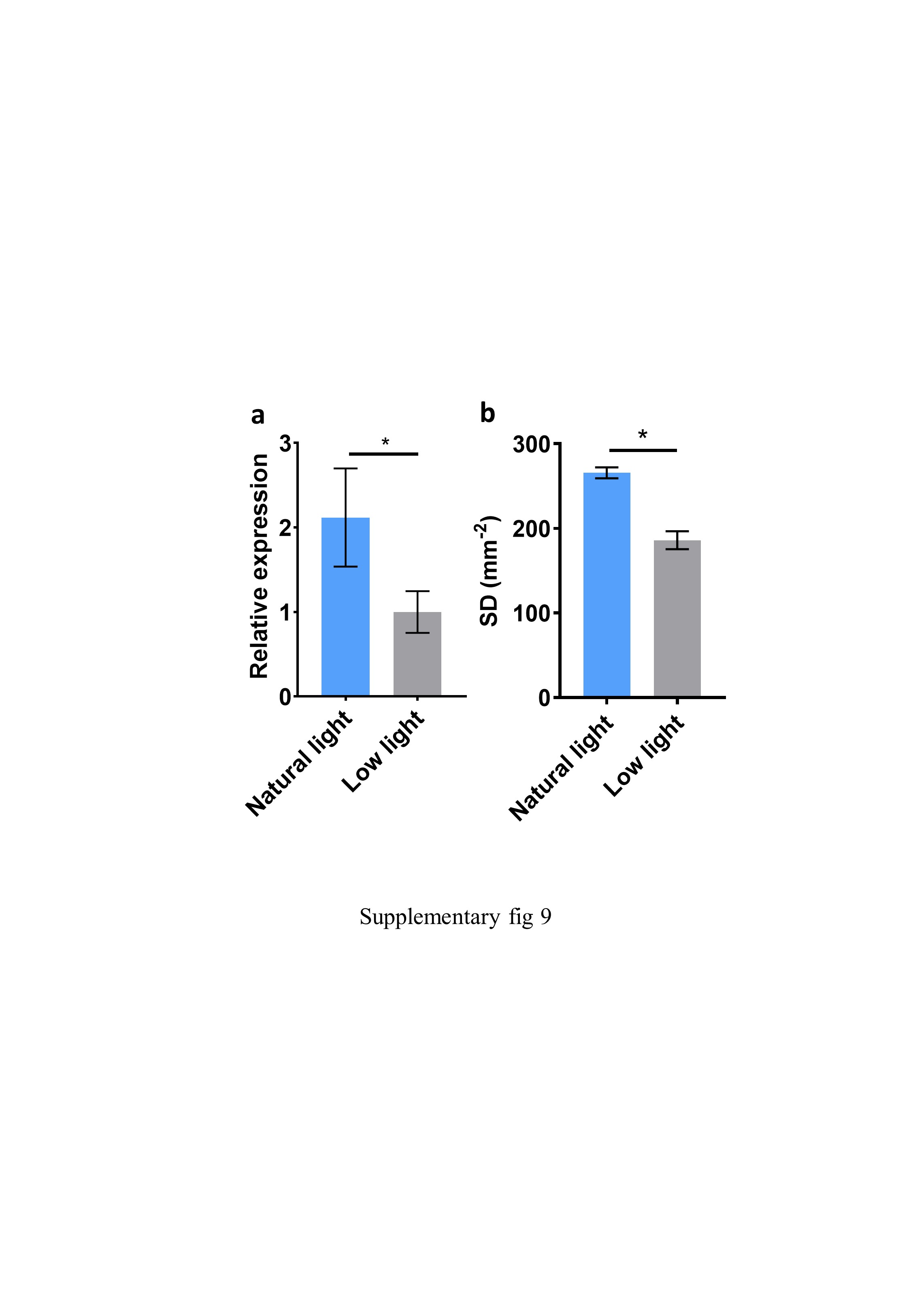

### supplementary fig 10

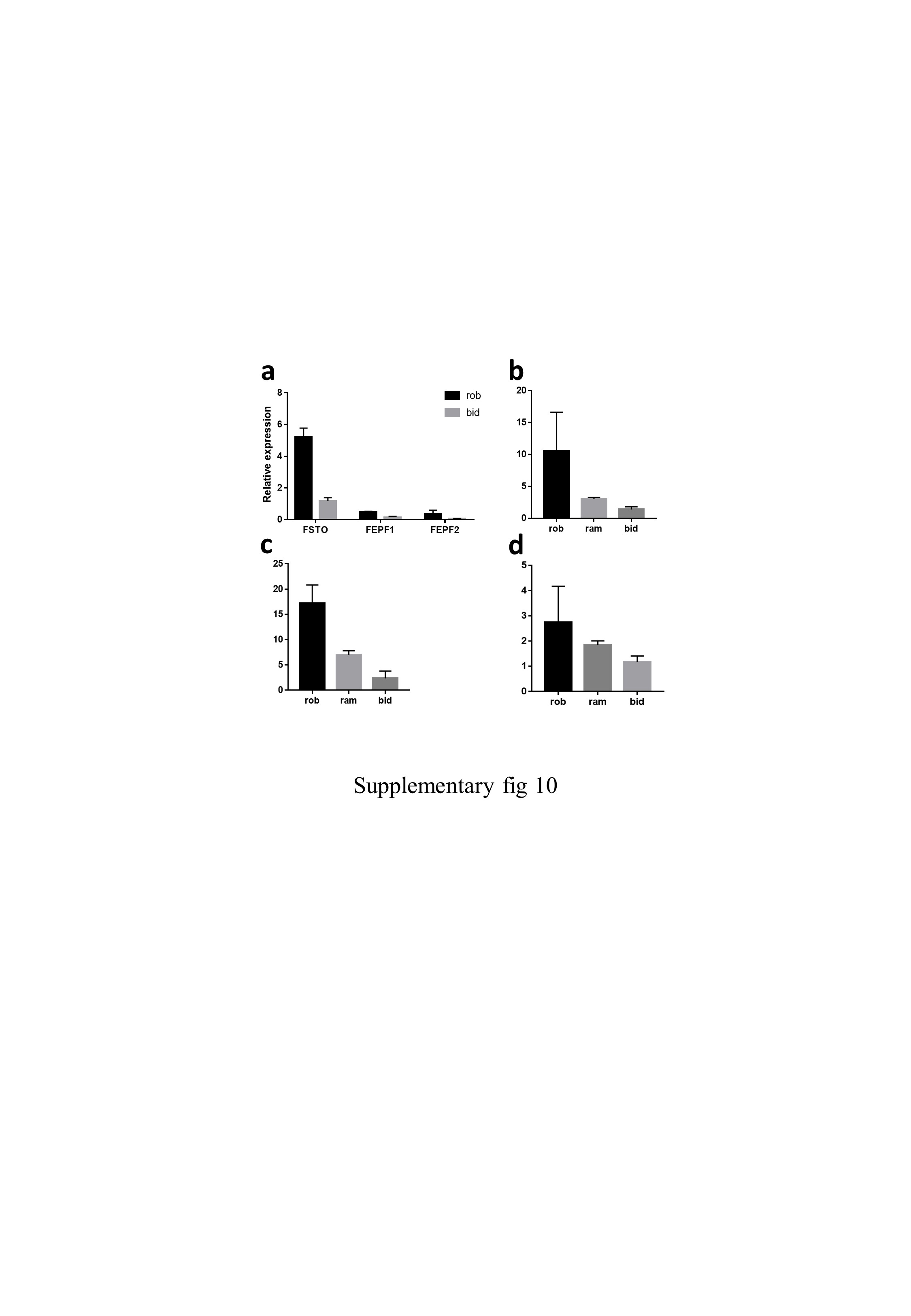

### supplementary fig 11

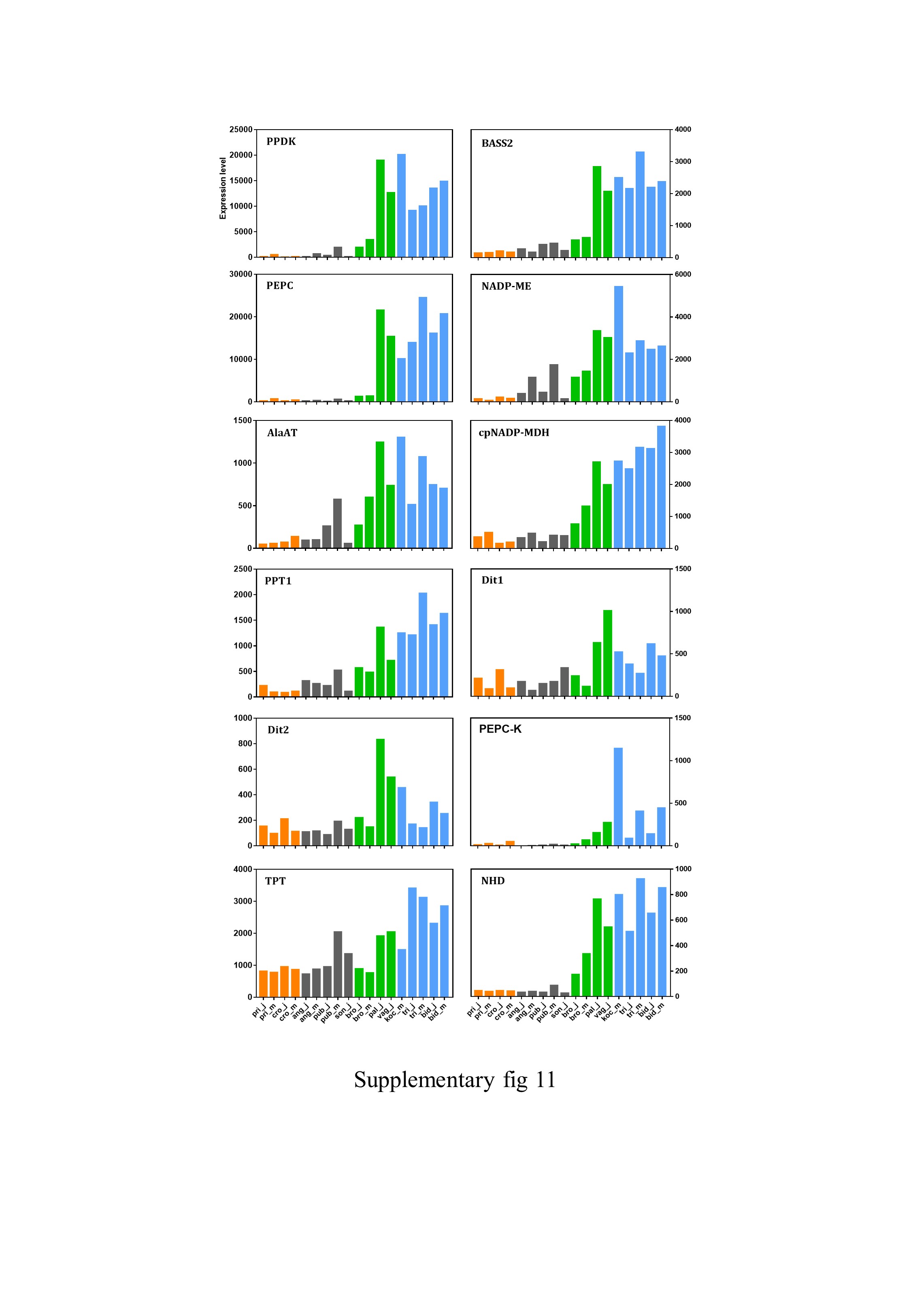

### supplementary fig 12

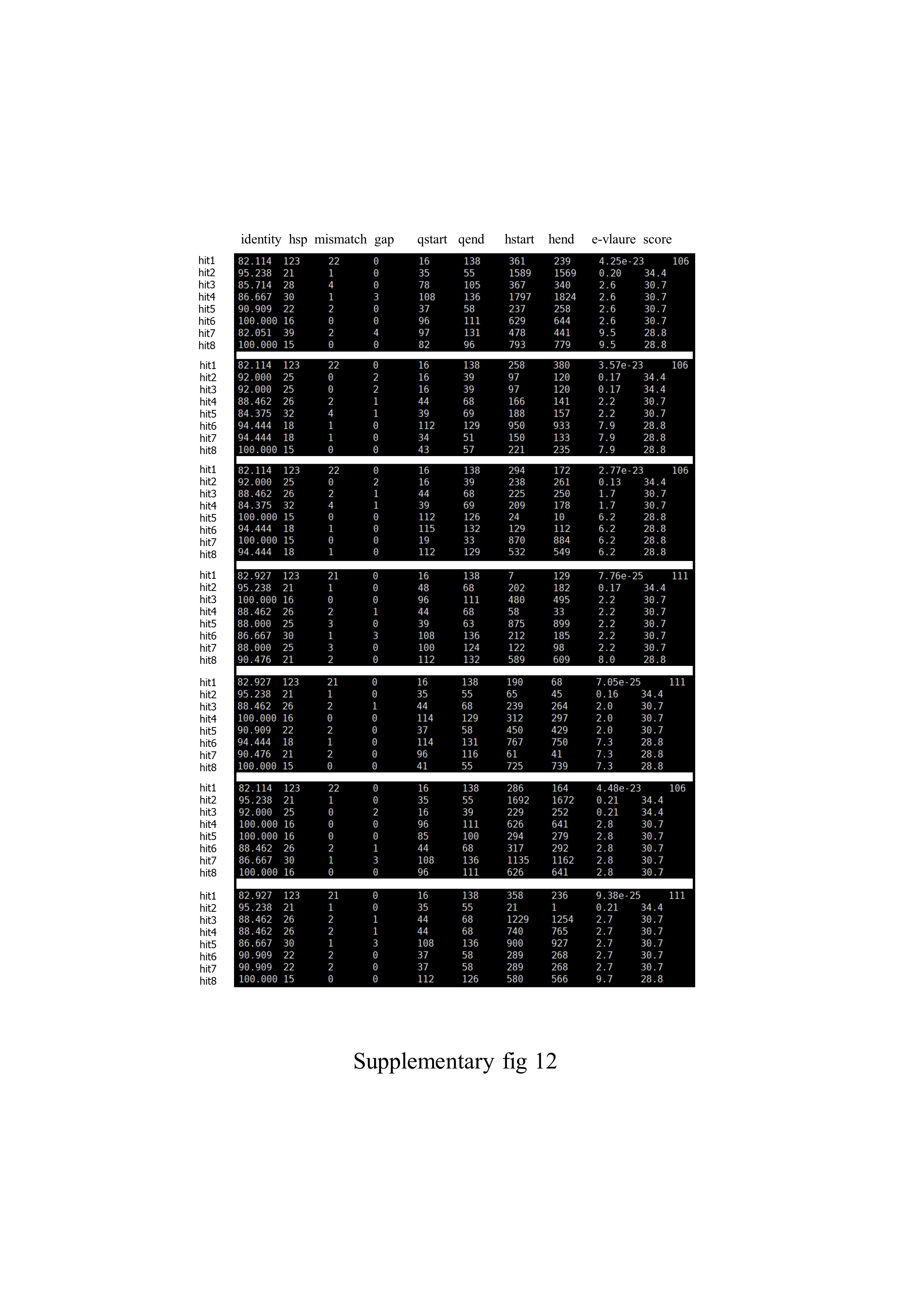

### supplementary fig 13

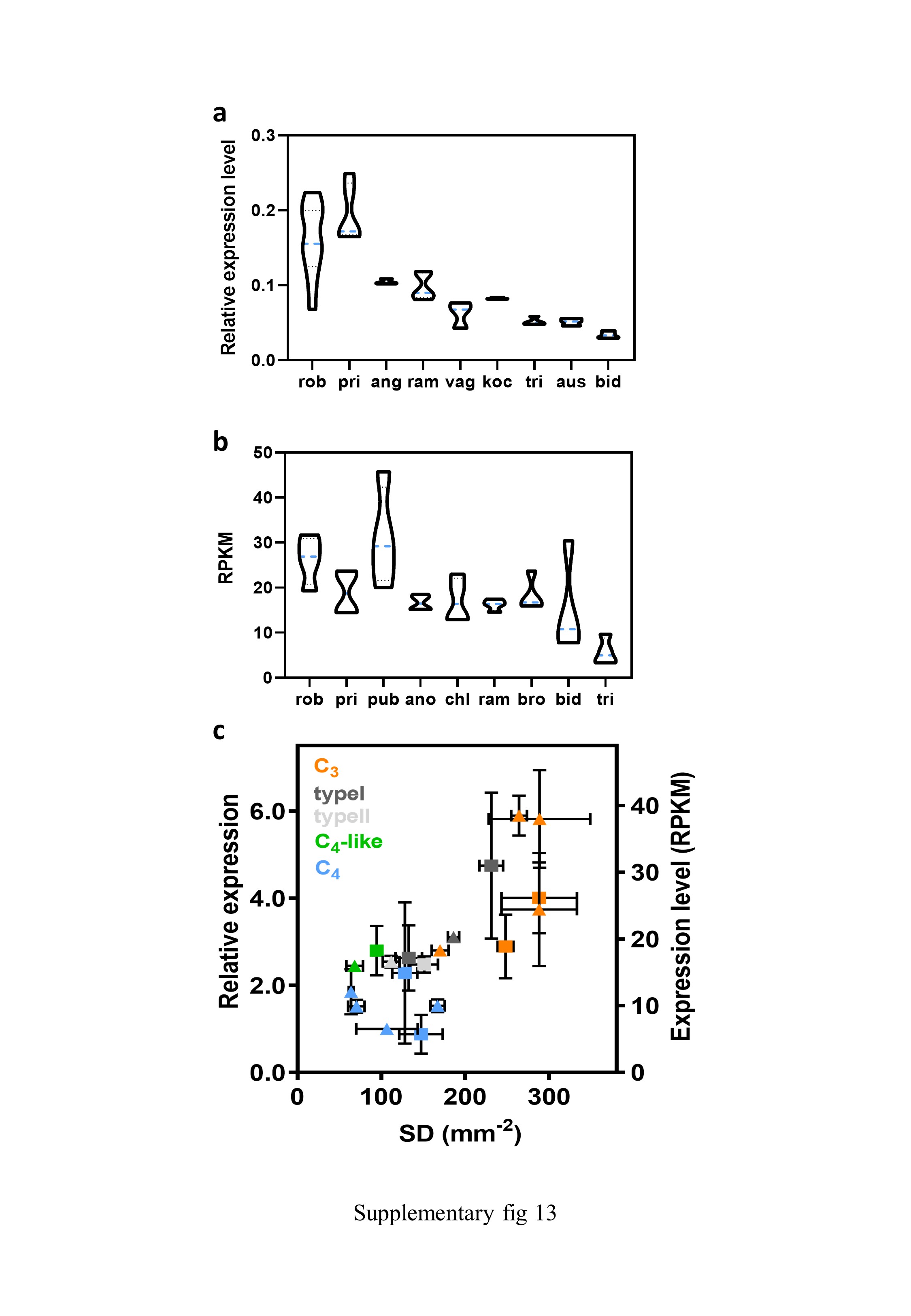

### supplementary fig 14

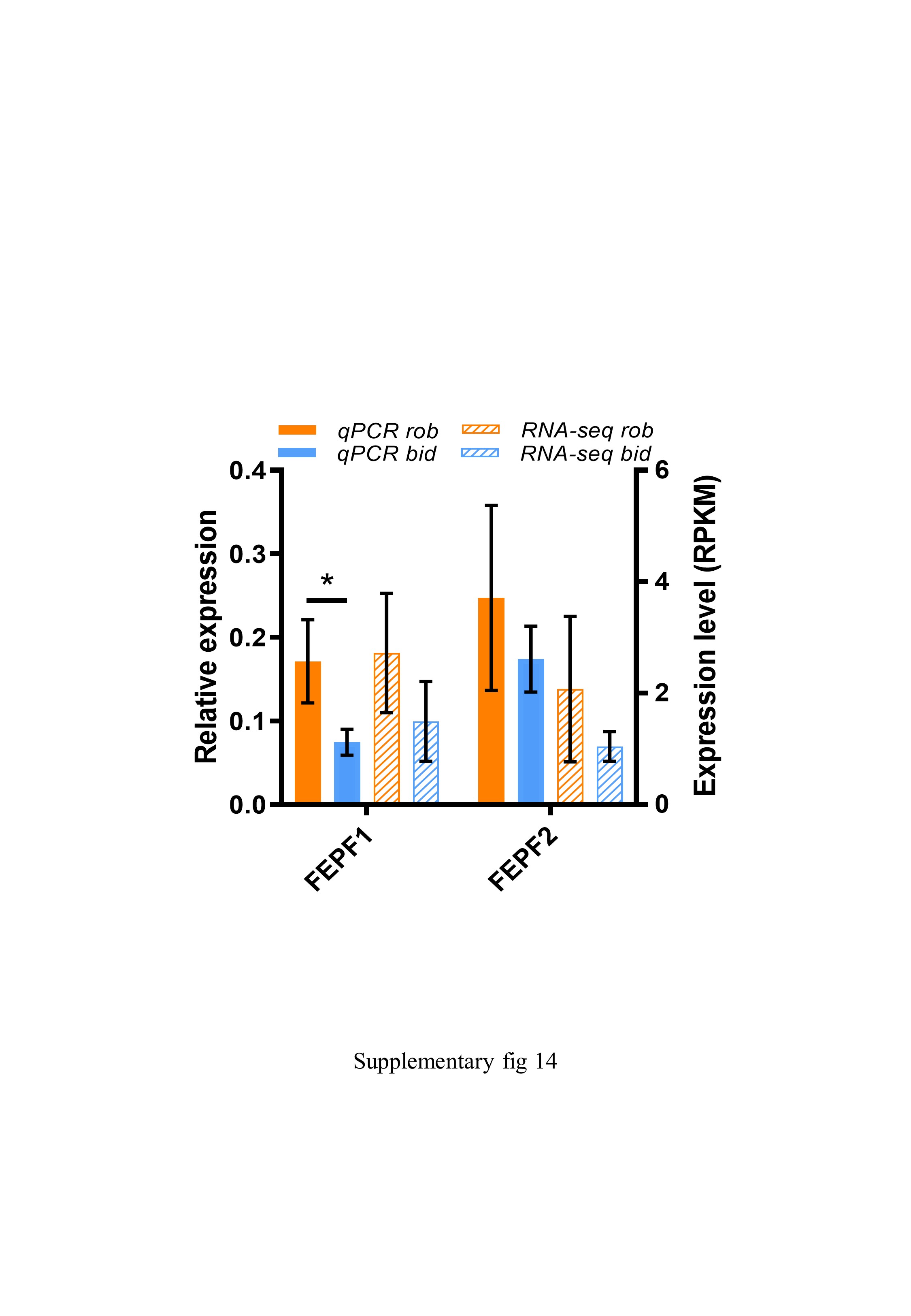

### supplementary fig 15

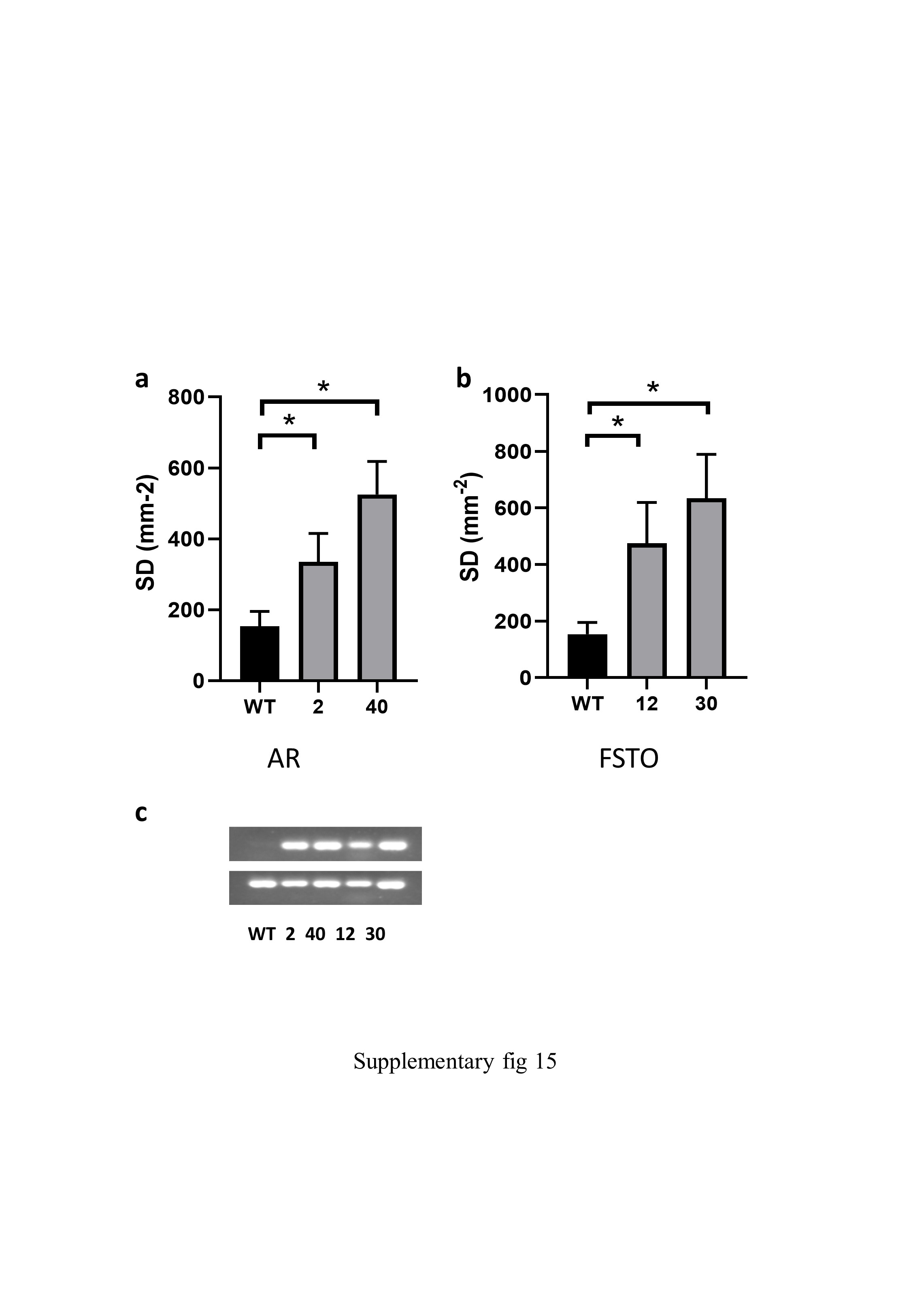

### supplementary fig 16

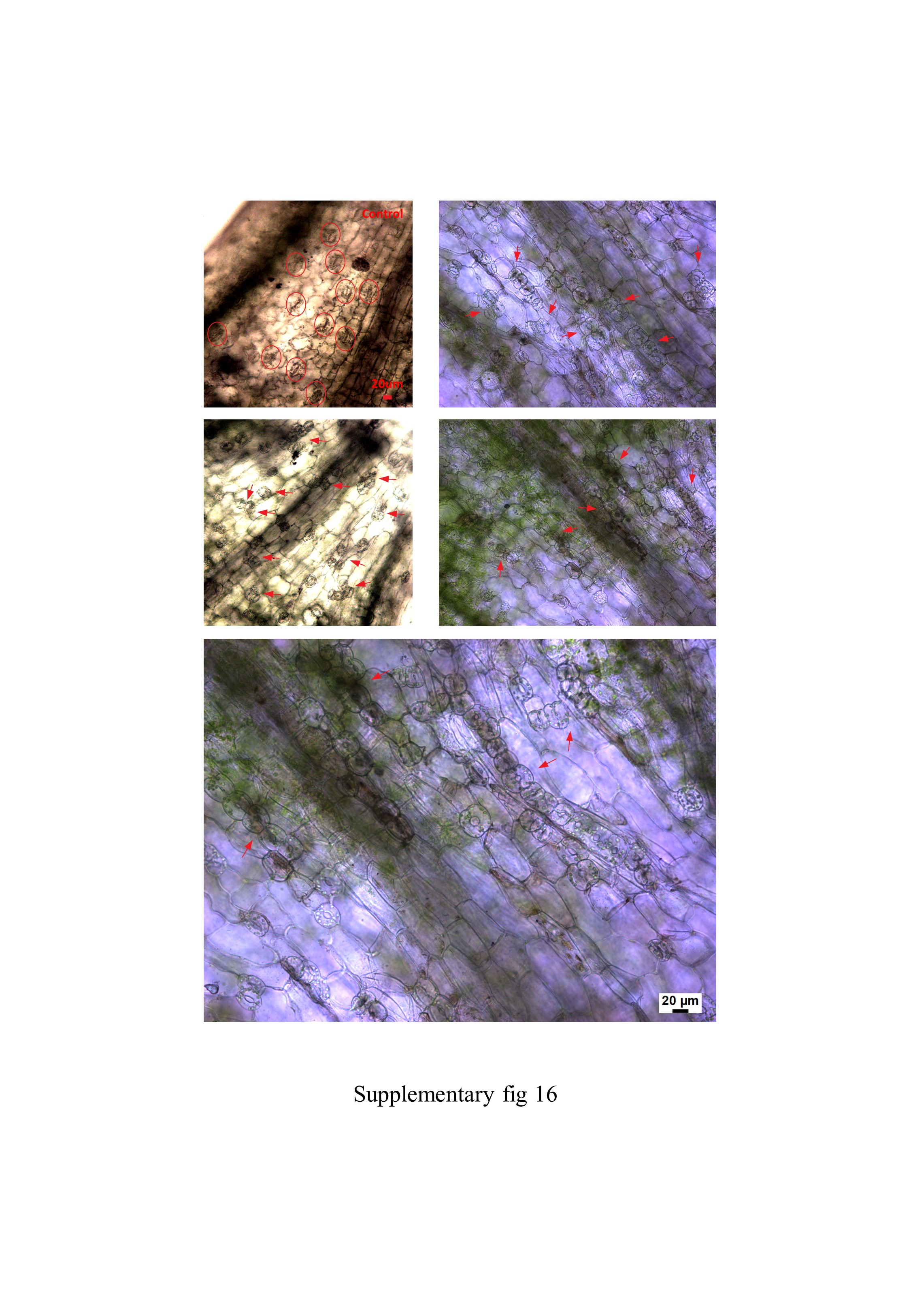

### supplementary fig 17

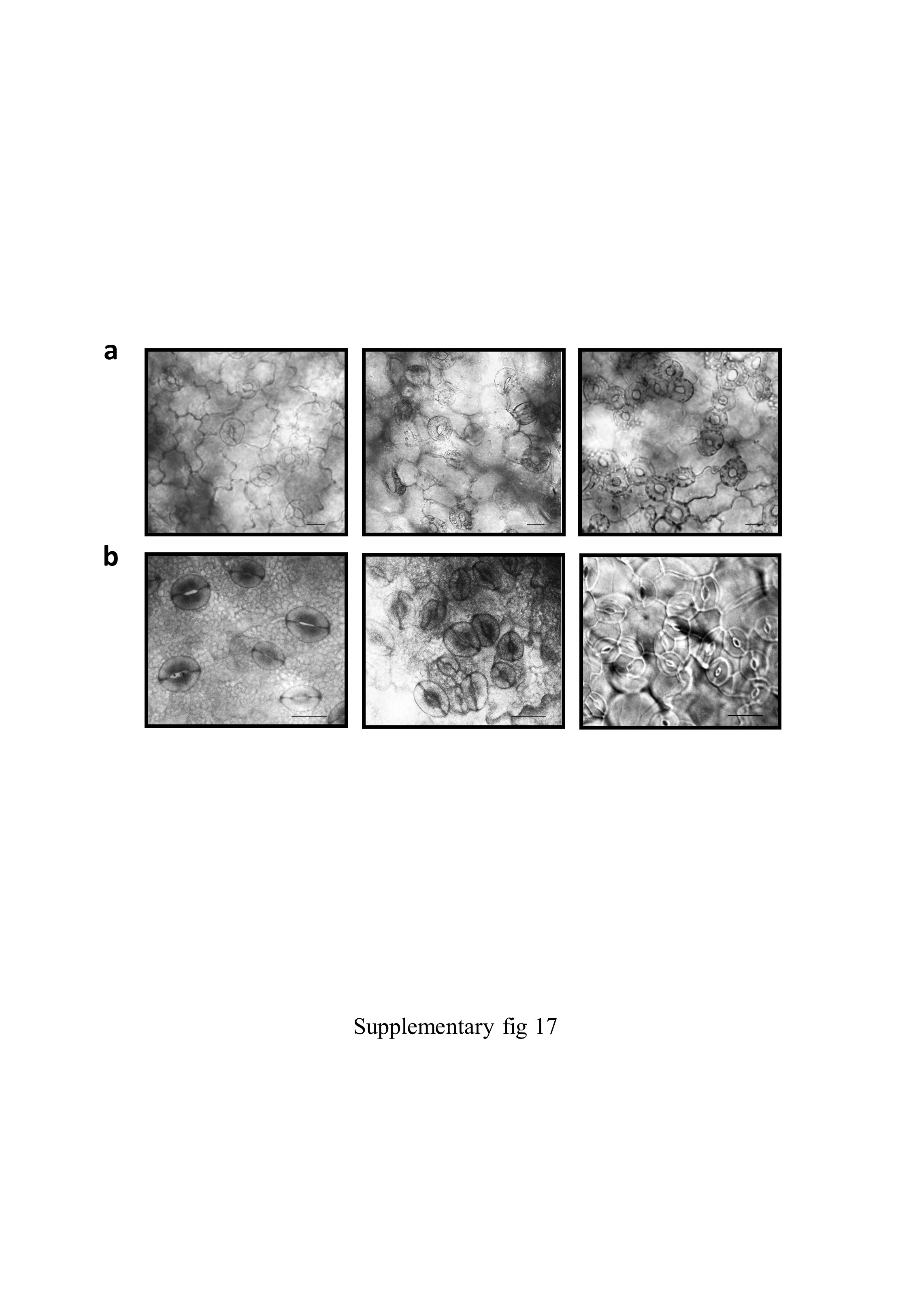

### supplementary fig 18

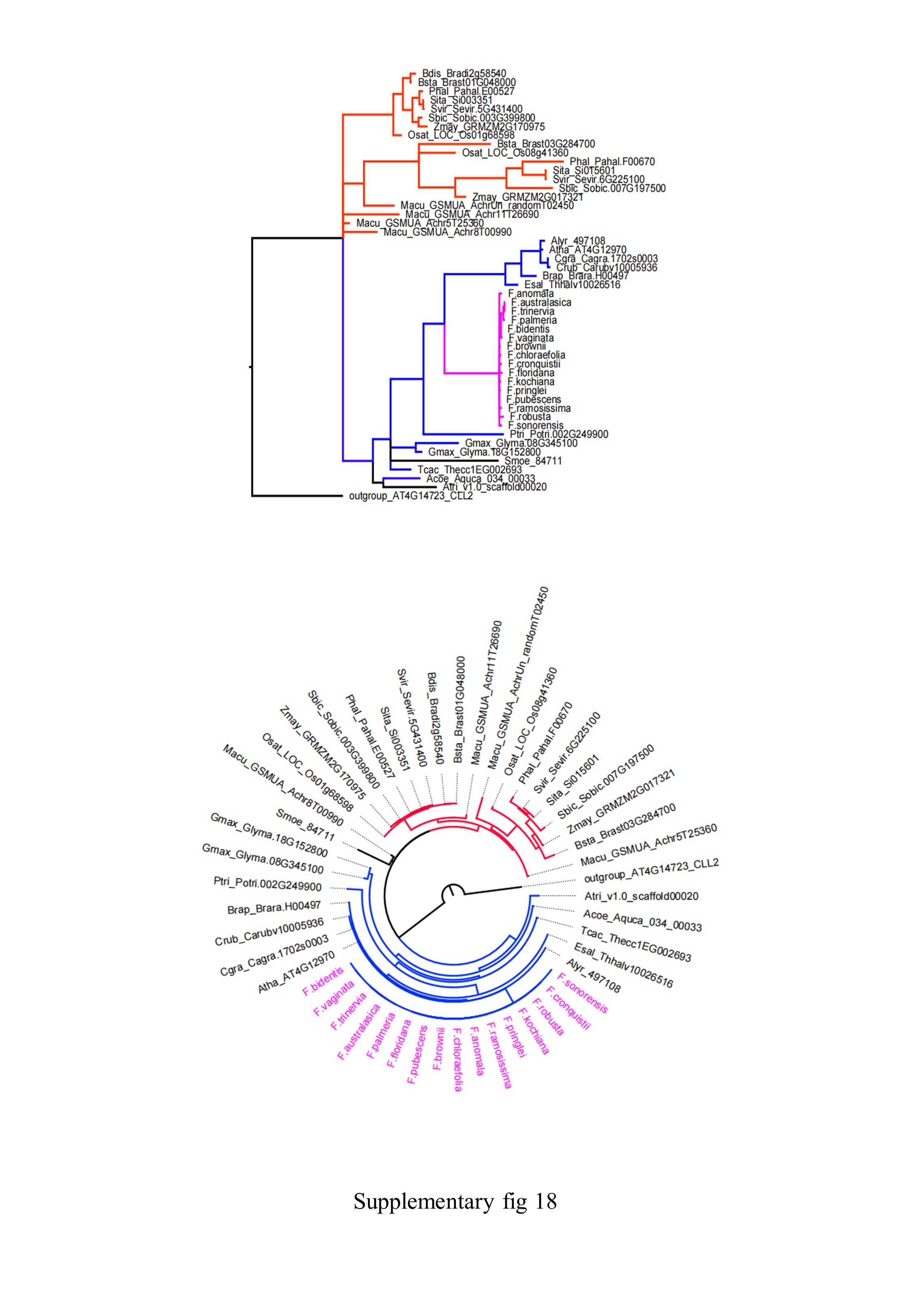

### supplementary fig 19

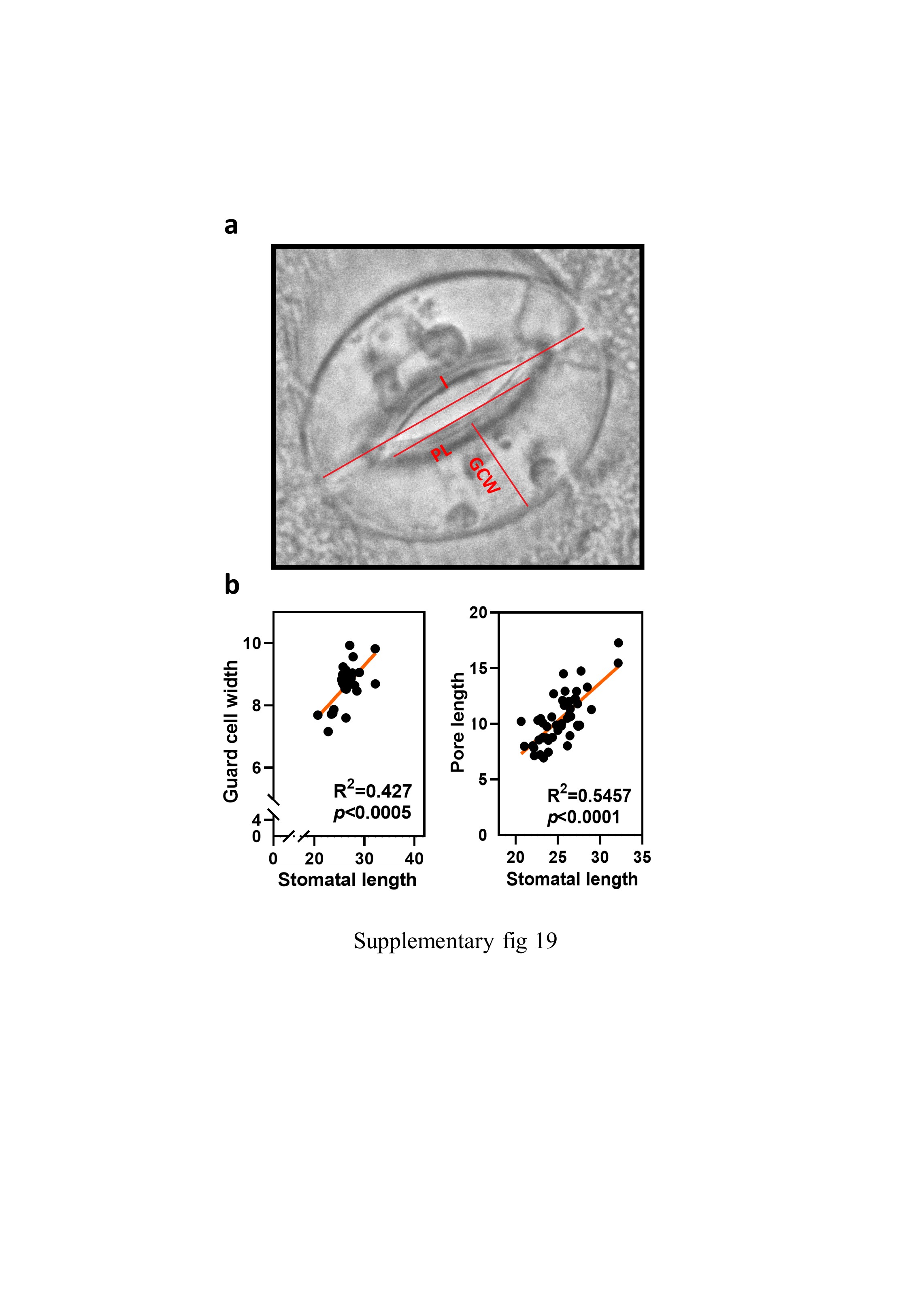

### supplementary fig 20

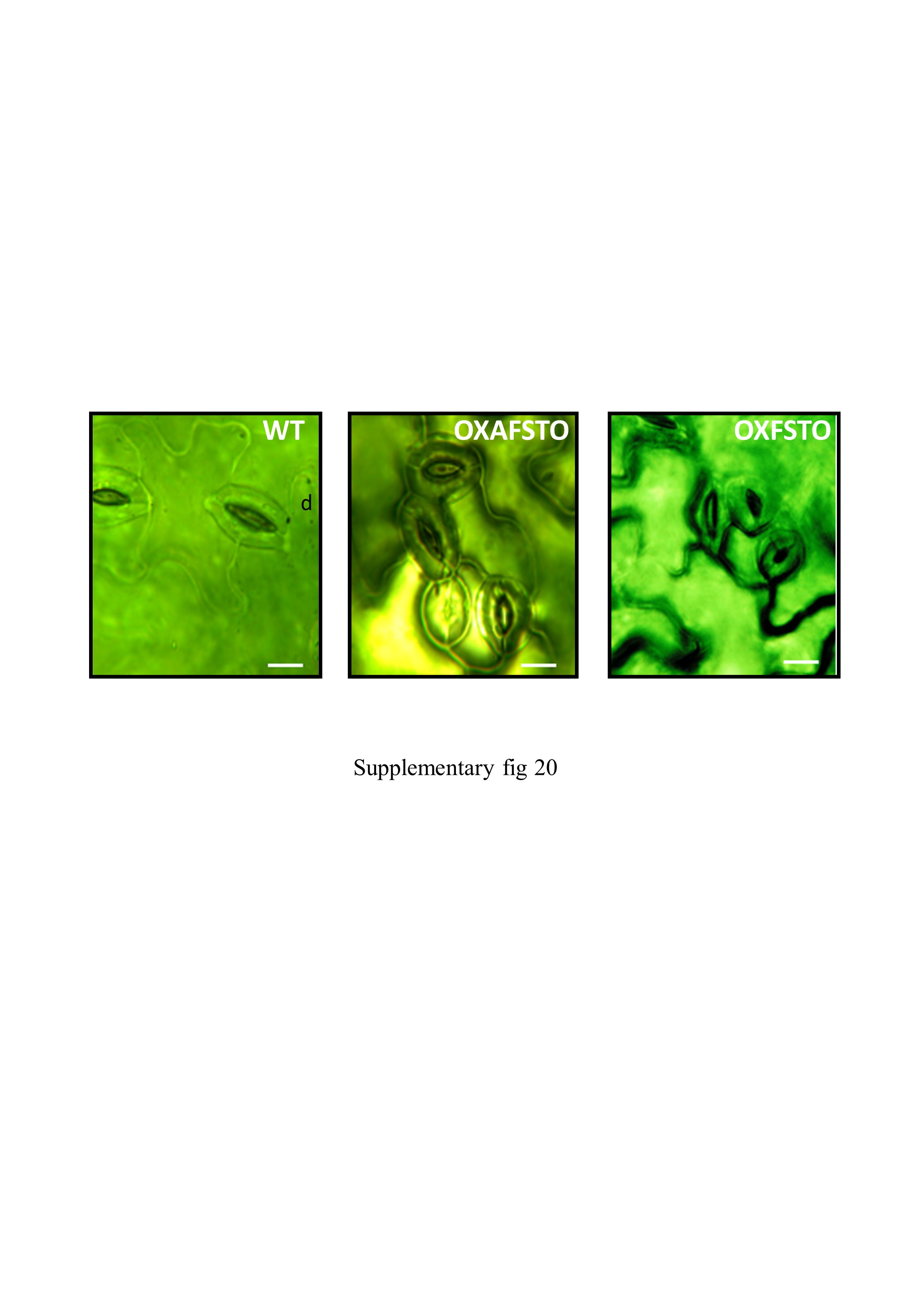

### supplementary table 1

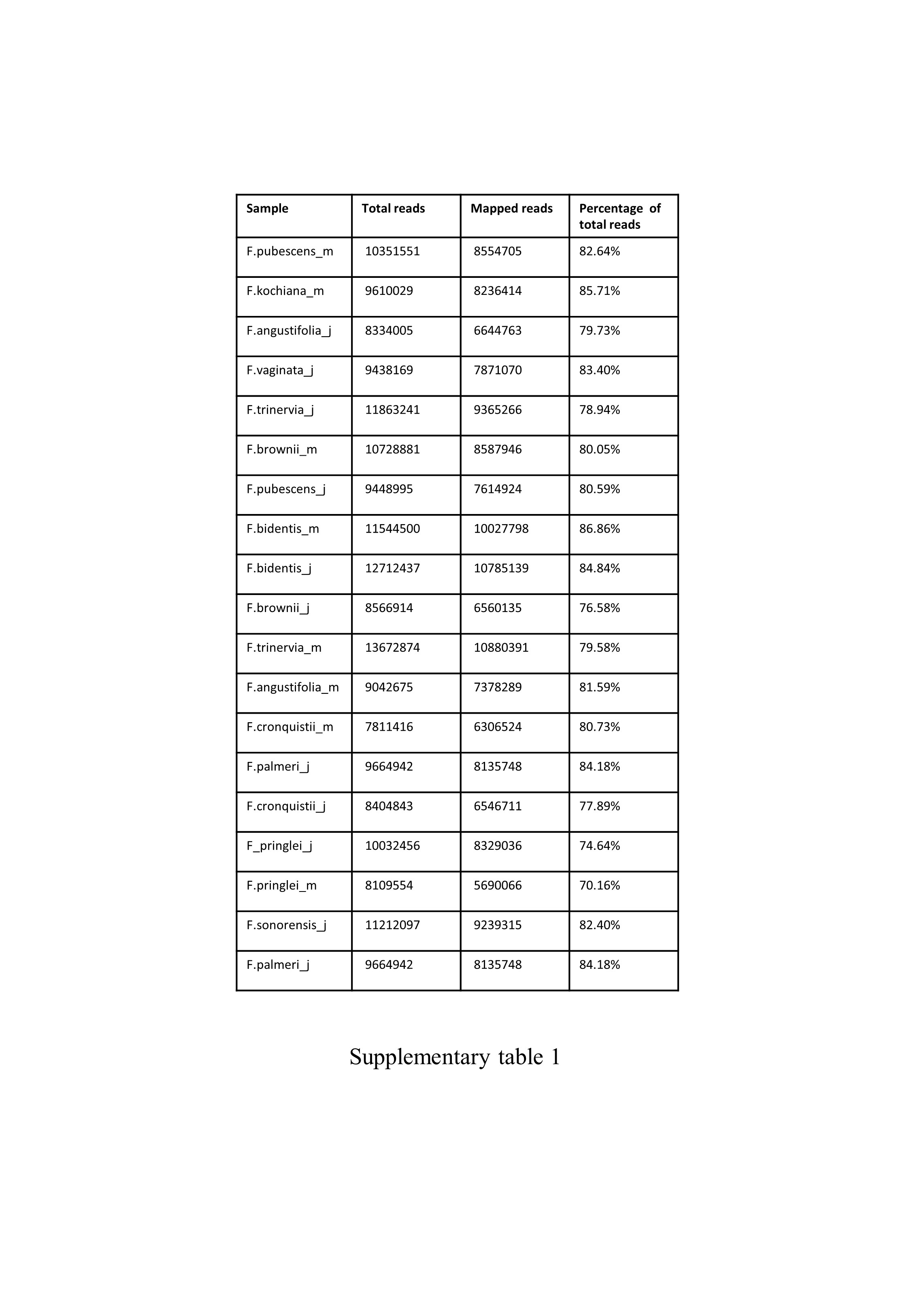

### supplementary table 2

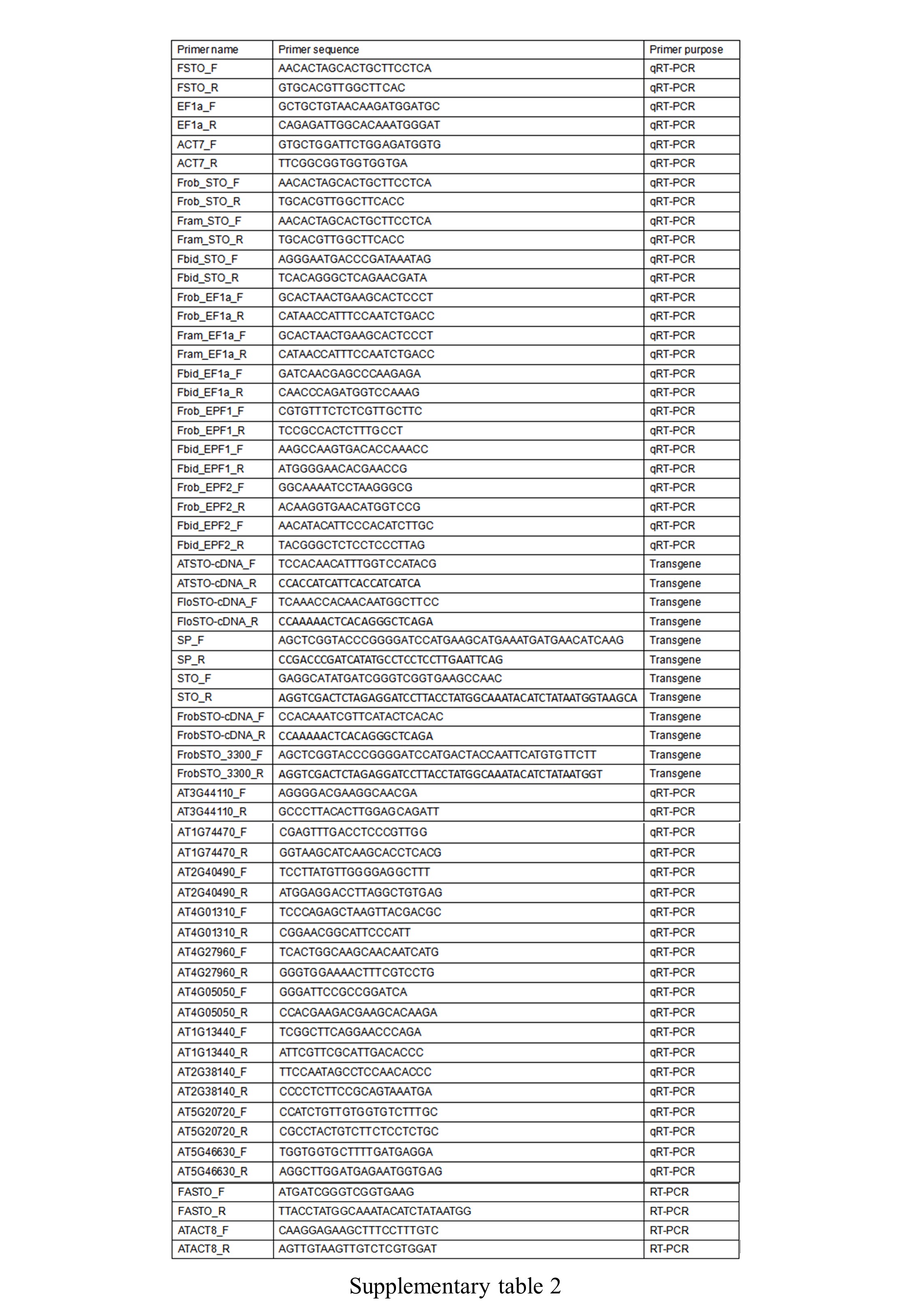

### supplementary table 3

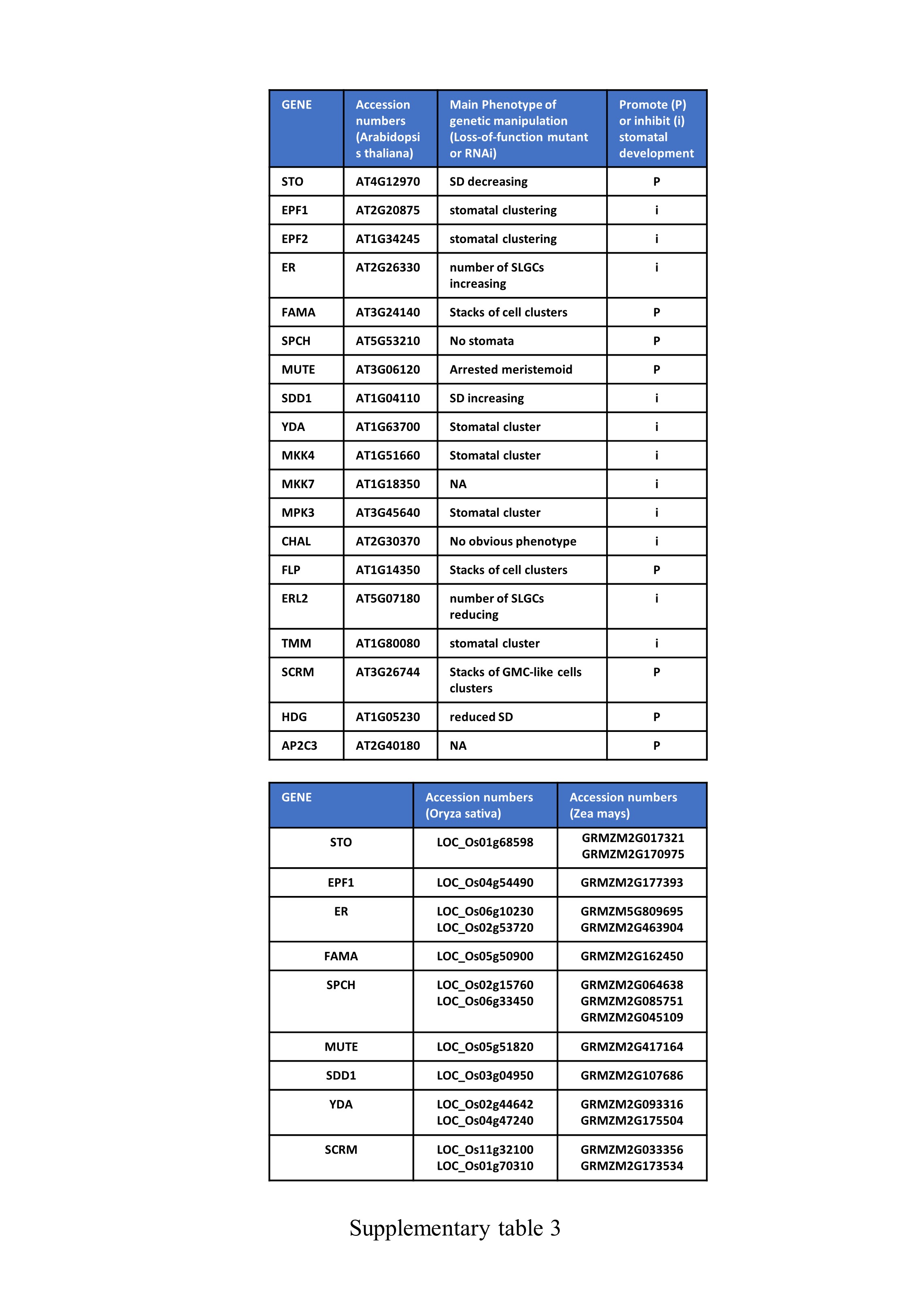

### supplementary table 4

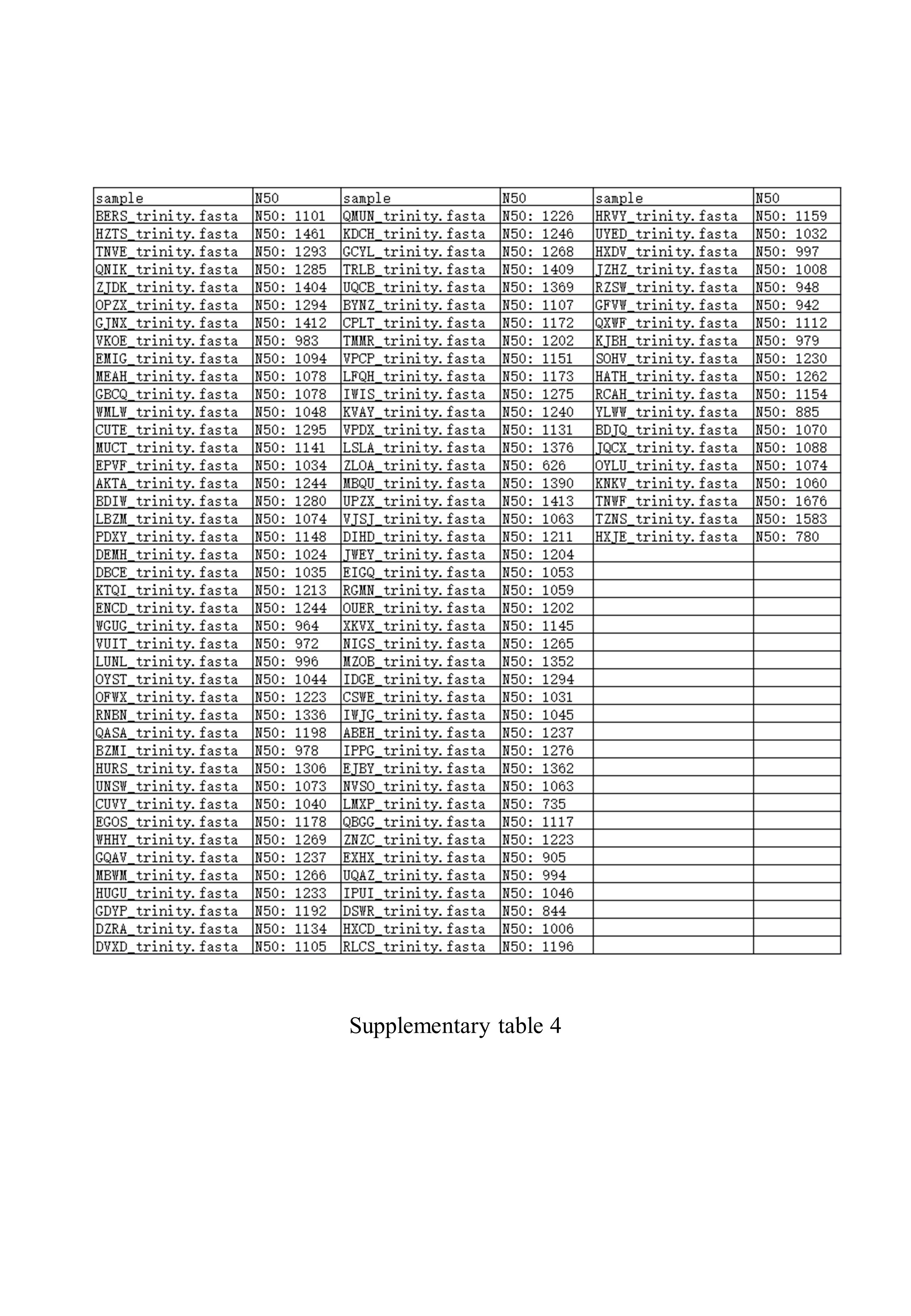

### supplementary table 5

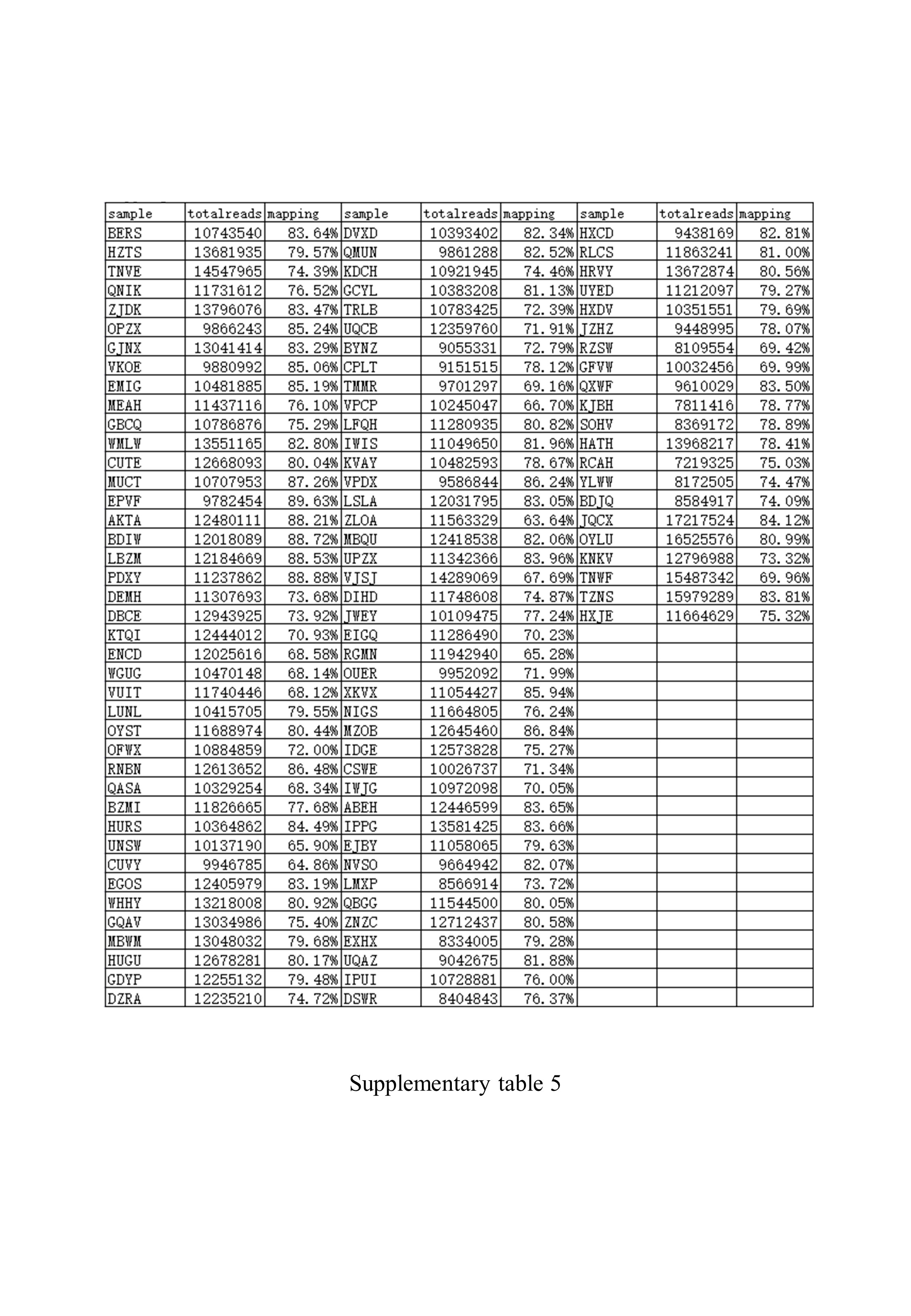
